# Time-resolved proteomic profiling of the ciliary Hedgehog response reveals that GPR161 and PKA undergo regulated co-exit from cilia

**DOI:** 10.1101/2020.07.29.225797

**Authors:** Elena A. May, Marian Kalocsay, Inès Galtier D’Auriac, Steven P. Gygi, Maxence V. Nachury, David U. Mick

## Abstract

The primary cilium is a signaling compartment that interprets Hedgehog signals through changes of its protein, lipid and second messenger compositions. Here, we combine proximity labeling of cilia with quantitative mass spectrometry to unbiasedly profile the time-dependent alterations of the ciliary proteome in response to Hedgehog. This approach correctly identifies the three factors known to undergo Hedgehog-regulated ciliary redistribution and reveals two such additional proteins. First, we find that a regulatory subunit of the cAMP-dependent protein kinase (PKA) rapidly exits cilia together with the G protein-coupled receptor GPR161 in response to Hedgehog; and we propose that the GPR161/PKA module senses and amplifies cAMP signals to modulate ciliary PKA activity. Second, we identify the putative phosphatase Paladin as a cell type-specific regulator of Hedgehog signaling that enters primary cilia upon pathway activation. The broad applicability of quantitative ciliary proteome profiling promises a rapid characterization of ciliopathies and their underlying signaling malfunctions.

## INTRODUCTION

The primary cilium is a solitary, microtubule-based protrusion of the cell that organizes developmental, sensory and homeostatic signaling pathways inside a functionally distinct compartment. Cilia defects cause multi-system pathologies named ciliopathies with symptoms including kidney cysts, retinal degeneration, obesity, brain malformations and skeletal anomalies (Reiter and Leroux, 2017; Hildebrandt et al., 2011). The vast range of symptoms underscores the broad physiological importance of cilium-based signaling. Our understanding of how cilia transduce signals is based in large part on studies of the developmental morphogen Hedgehog (Hh) (Gigante and Caspary, 2020; Anvarian et al., 2019; Kong et al., 2019). In vertebrates, Hh signaling is strictly dependent on an intact primary cilium. The core Hh machinery comprises the Hh receptor Patched (PTCH1), the G-protein coupled receptors (GPCR) GPR161 and Smoothened (SMO), and the GLI transcription factors, all of which dynamically localize to primary cilia in response to Hh (Fig. 1A). PTCH1 and GPR161, two molecules that restrain Hh pathway activation inside cilia in unstimulated cells, undergo ciliary exit upon pathway stimulation while the central pathway activator SMO becomes enriched inside cilia when activated. It has been proposed that PTCH1 pumps a lipidic activator of SMO out of the ciliary membrane and that the PTCH1 lipid extruder activity is directly suppressed upon liganding Hh. Ciliary exit of PTCH1 further reduces the inhibitory effect exerted by PTCH1 on SMO. Downstream of SMO and GPR161 lies the cAMP-dependent protein kinase (PKA) which phosphorylates GLI2 and GLI3 and commits them to processing into transcriptional repressors. It is now believed that it is the PKA activity within cilia that is important for transduction of Hh signals (Kong et al., 2019; Gigante and Caspary, 2020). SMO blocks PKA activity, either directly (Arveseth et al., 2020) or via its Gα_i_-mediated inhibition of adenylyl cyclases (Riobo, 2014) and GPR161 increases PKA activity within cilia via its tonic activation of adenylyl cyclases though Gα_s_. By triggering the ciliary exit of GPR161, the accumulation of activated SMO in cilia thus results in a drastic decrease in ciliary PKA activity (Mukhopadhyay et al., 2013; Pusapati et al., 2018a), and a switch in the processing of the transcription factors GLI2/3 from repressor to activator forms. The dynamic redistribution of signaling molecules thus plays an integral part in the transduction of Hh signals inside the cilium. Despite the recognized importance of dynamic ciliary localization in the Hh response, the extent of ciliary proteome remodeling during Hh signaling remains unknown and several of the key steps remain incompletely characterized. For example, the exact mechanism of how ciliary SMO triggers the exit of GPR161 from cilia remains to be determined.

**Figure 1.**
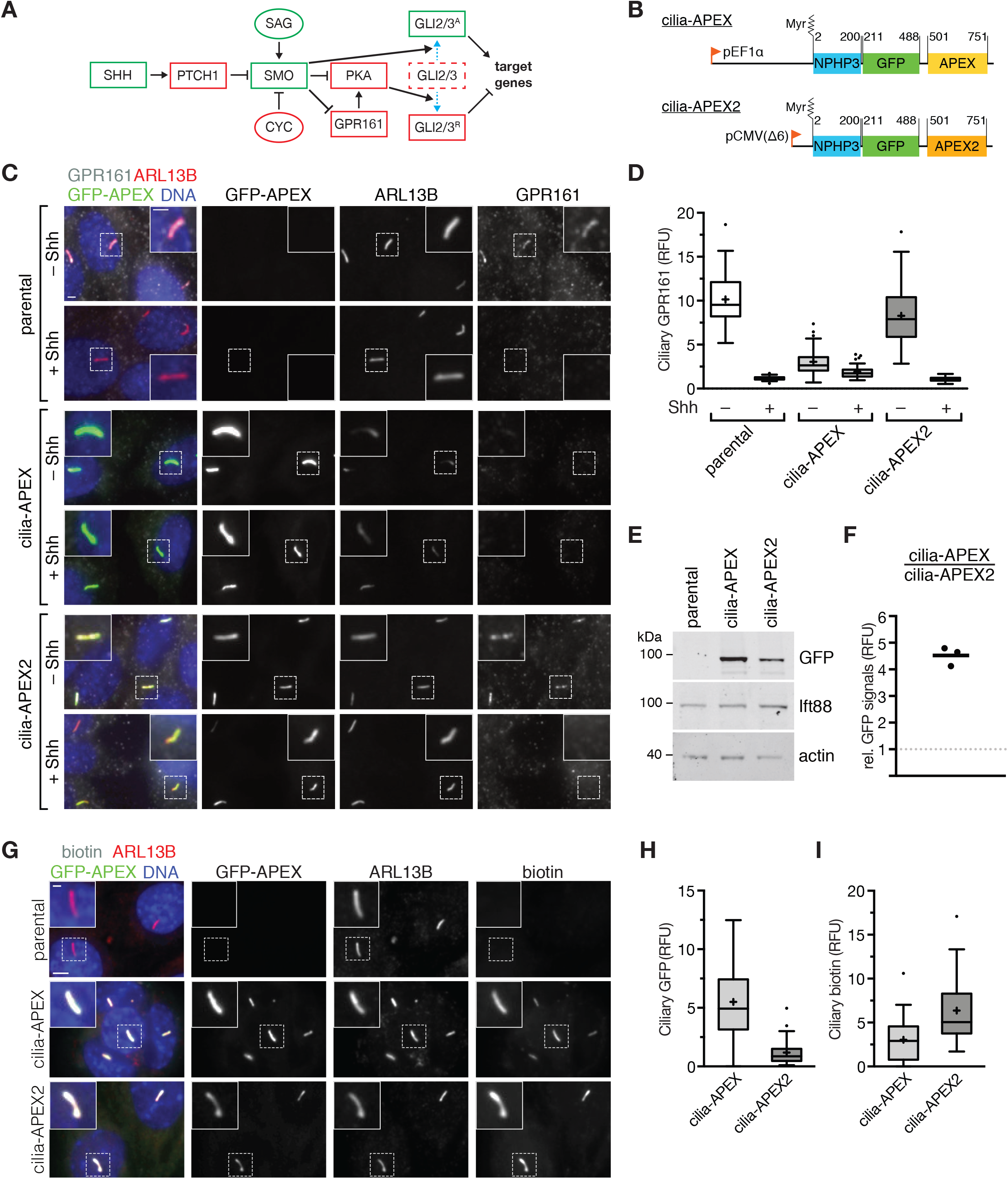
An IMCD3 cell line expressing cilia-APEX2 at low levels efficiently labels proteins of the primary cilium without disrupting ciliary localization of Hh signaling components. **(A)** Schematic overview of the key Sonic Hedgehog signaling components. Positive and negative regulators are indicated in green and red boxes, respectively. Pharmacological agents used in this study are displayed in ovals. Smoothened agonist (SAG) activates the pathway, while cyclopamine (CYC) inhibits the pathway at the level of SMO. **(B)** Diagram of the *cilia-APEX* and *cilia-APEX2* transgenes. Both contain a cilia-targeting signal based on the N-terminus of NPHP3, which is myristoylated at a glycine residue at position 2 (Myr), followed by a GFP moiety. Numbers indicate amino acid positions. cilia-APEX is expressed from an EF1a promoter (pEF1a), cilia-APEX2 from a truncated CMV promoter (pCMV(Δ6)) for reduced expression. **(C)** Immunofluorescence of parental IMCD3 cells and of stable clones expressing cilia-APEX or cilia-APEX2. Cells were serum-starved for 24 h in the presence or absence of Sonic Hedgehog-conditioned medium (Shh), fixed and stained for GPR161 (white) and ARL13B (red) using specific antibodies. Cilia-APEX and cilia-APEX2 were visualized via the intrinsic fluorescence of GFP (green). DNA was stained with DAPI (blue). Representative micrographs are shown for each condition. **(D)** Box plots showing the relative GPR161 fluorescence normalized to ARL13B in the primary cilium of parental, cilia-APEX and cilia-APEX2 cell lines after Shh treatment as in (**A**). 50 cilia were analyzed for each condition (n = 50). In these and all subsequent box plots, crosses indicate mean values, whiskers indicate values within 1.5x interquartile range and dots represent outliers. **(E)** Cell lysates of parental, cilia-APEX and cilia-APEX2 cell lines were resolved by SDS-PAGE and analyzed by quantitative immunoblotting using indicated antibodies. APEX fusion proteins were detected using anti-GFP antibodies. **(F)** Dot plot showing the ratio of total GFP protein detected in the cilia-APEX relative to the cilia-APEX2 cell line (n = 3). **(G)** Parental, cilia-APEX and cilia-APEX2 IMCD3 cells were subjected to APEX labeling, fixed and stained for ARL13B (red) and biotin (white). Cilia-APEX and cilia-APEX2 were visualized via the intrinsic fluorescence of GFP (green). DNA was stained with DAPI (blue). Representative micrographs are shown. **(H** and **I)** Box plots showing background-subtracted intensities of GFP (**I**) and biotin (**J**) signals in the primary cilium from images as in (E). 30 cilia were analyzed for each condition (n = 30). Scale bars = 2 μm in all panels.

An unbiased, systematic description of the mammalian primary cilia proteome has proven challenging because the isolation of mammalian cilia remains fraught with severe limitations (Ishikawa et al., 2012). To overcome this technical barrier, we previously established a proximity labeling-based method using cilia-localized ascorbate peroxidase (cilia-APEX) (Mick et al., 2015). Cilia-targeted APEX fusion proteins allow labeling of ciliary components with biotin-derivatives that are readily enriched by streptavidin chromatography (Mick et al., 2015; Kohli et al., 2017). Proteomics of cilia using cilia-APEX has contributed to our molecular understanding of Hh signaling (Mick et al., 2015), of extracellular vesicles shedding from cilia (Nager et al., 2017) and of regulated ciliary GPCR trafficking (Shinde et al., 2020). While APEX approaches in cilia have identified several ciliary signaling proteins (Mick et al., 2015; Kohli et al., 2017), they failed to robustly detect signaling proteins of high importance and low abundance, such as PTCH1, GPR161 and SMO. The scarcity of these factors in cilia combined with the high hydrophobicity of these multi-pass membrane proteins makes mass spectrometric analysis a considerable challenge. The limited sensitivity of cilia-APEX thus hampers a systematic and quantitative assessment of the proteomic changes in cilia in response to signals and other perturbations.

Here, we profile the changes of the ciliary proteome during Hh signaling in a systematic and time-resolved manner by combining improvements in the cilia-APEX2 proximity labeling scheme with state-of-the-art quantitative mass spectrometry using tandem-mass-tags (TMTs) (Paek et al., 2017). These technical advances enabled us to uncover the extent of the ciliary proteome remodeling in response to Hh ligand and gain novel mechanistic insights into Hh signaling.

## RESULTS AND DISCUSSION

### The cilia-APEX2 expression system limits perturbations of ciliary dynamics

To investigate why cilia-APEX had failed to identify ciliary membrane proteins of the Hh pathway, we surveyed the ciliary accumulation of these Hh signaling molecules by immunofluorescence microscopy in the inner medullary collecting duct (IMCD3) cell line stably expressing the cilia-APEX transgene (Fig. 1B). Surprisingly, while endogenous GPR161 was readily detected in cilia of parental IMCD3 cells, ciliary GPR161 was nearly undetectable in IMCD3-[cilia-APEX] cells (Figs. 1C and 1D). The relatively high level of cilia-APEX expression required to support efficient biotinylation may have altered ciliary composition. Even though a weak promoter (pEF1α) was used to stably express cilia-APEX and expression was limited by integration at a defined genomic locus, we previously found that pEF1α-driven expression of a ciliary GPCR from the same locus produced nearly 50,000 molecules per cilium and led to a drastic lengthening of cilia (Ye et al., 2018).

We thus reduced expression levels of the transgene by switching to the truncated CMV(Δ6) promoter (Morita et al., 2012), a considerably weaker promoter than pEF1α (Ye et al., 2018). To maintain robust biotinylation levels, we leveraged APEX2, an APEX variant with increased turnover rates (Lam et al., 2015). As for the original cilia-APEX, ciliary targeting was carried out by the N-terminus of Nphp3 (Wright et al., 2011; Nakata et al., 2012), and the presence of a GFP moiety in both cilia-APEX and cilia-APEX2 allowed us to compare their relative abundance (Fig. 1B). The total levels of cilia-APEX2 were reduced nearly 5-fold compared to cilia-APEX in the respective cell lines (Figs. 1E and 1F) while the abundance of the ciliary protein IFT88 was not affected (Figs. 1E and S1A). Congruently, the ciliary intensity of cilia-APEX2 was reduced about 5-fold compared to cilia-APEX (Figs. 1G and 1H). Despite the drastic reduction in ciliary abundance of cilia-APEX2 compared to cilia-APEX, ciliary biotinylation efficiency in the presence of the APEX substrates biotin tyramide and H_2_O_2_ nearly doubled in cilia-APEX2 compared to cilia-APEX (Figs. 1G and 1I). Importantly, the ciliary abundance of GPR161 was nearly indistinguishable between parental and cilia-APEX2 cells (Figs. 1C and 1D). Furthermore, GPR161 was efficiently removed from cilia in response to Hh pathway activation by Sonic Hedgehog (Shh) in the cilia-APEX2 cell line, confirming that the IMCD3-[cilia-APEX2] cell line recapitulates the physiological ciliary dynamics of the Hh signaling molecules in response to Hh pathway activation. Together, these results predict that cilia-APEX2 improves sensitivity of ciliary proteomics while minimizing perturbations of ciliary protein dynamics.

### Tandem mass tag analyses of cilia-APEX2 samples extend coverage of the ciliary proteome

We next sought to compare the performance of cilia-APEX2 to that of cilia-APEX in quantitative mass spectrometric analysis. After labeling, biotinylated proteins were isolated via streptavidin chromatography and quantified by tandem mass tag labeling and mass-spectrometry. In addition to proteins biotinylated by the APEX enzyme, we expected streptavidin enrichment of several endogenously biotinylated proteins as well as proteins biotinylated by endogenous peroxidases during the labeling reaction. To discriminate proteins directly labeled by cilia-APEX2 from other biotinylated proteins, we leveraged several controls. First, we introduced a single amino acid substitution in the myristoylation site of the Nphp3-based ciliary targeting signal to abolish cilia targeting (Wright et al., 2011; Nakata et al., 2012), thus establishing control-APEX2. Expressed from the same locus and promoter as cilia-APEX2, control-APEX2 identifies the proteins labeled by the cilia-APEX2 molecules that did not reach the cilium (e.g. biogenesis intermediates). Secondly, to further control for non-ciliary proteins, in particular membrane proteins that may not be labeled by control-APEX2 (a soluble protein), we genetically ablated cilia in the cilia-APEX2 cell line by disrupting the centriole distal appendage protein CEP164 using CRISPR/Cas9-mediated genome editing (Fig. S1B). Basal bodies cannot dock to the plasma membrane in the absence of CEP164, thus precluding cilium assembly without effecting other cellular processes (Daly et al., 2016; Tanos et al., 2013). *Cep164^−/−^* IMCD3-[cilia-APEX2] cells lack cilia (Fig. S1C-D) while retaining expression of the identical fusion protein that is used to label ciliary proteins in wild-type cells.

We conducted synchronous precursor selection MS/MS/MS (MS^3^) analyses of APEX-labeled samples from WT cilia-APEX2, *Cep164^−/−^* cilia-APEX2, and control-APEX2 cell lines in triplicate after streptavidin capture and tandem mass tag (TMT) labeling of tryptic peptides (Fig. 2A). The TMT isobaric tagging approach enables precise and reproducible quantification of the relative protein abundances in different samples (Liu et al., 2020; Paek et al., 2017; Li et al., 2020). Hierarchical clustering of each protein’s relative abundance in the ten samples analyzed within one multiplex experiment demonstrates the high reproducibility of the triplicate experiment (Fig. S1E). Proteins that are highly enriched in the cilia-APEX2 data set compared to the controls form two clusters of candidate ciliary proteins (Fig. S1E and Fig. 2B) while non-ciliary proteins fall into separate clusters (Fig. S1F). We defined the cilia-APEX2 proteome via statistical analyses of the relative enrichment between the cilia-APEX2 samples and the controls (Fig. 2C and 2D). To be included in the cilia-APEX2 proteome, proteins had to fulfill four criteria (Figs. 2C-D, blue dots): greater than 2-fold enrichments in the cilia-APEX2 samples over control-APEX2 samples and over the *Cep164^−/−^* cilia-APEX2 samples (TMT ratio > 2.0), and statistically significant enrichments in the cilia-APEX2 samples vs. the control-APEX2 samples and the *Cep164^−/−^* cilia-APEX2 samples (p-value < 0.05). In addition, proteins that fulfilled only 3 out of 4 criteria (Figs. 2C-D, black dots) were included if they were close to matching the fourth criterion (TMT ratio > 1.5, or p-value < 0.1). This set of criteria resulted in the inclusion of 203 proteins (Table S1) and represents a compromise between inclusion of false positives and exclusion of false negatives (Fig. 2C-D). It should be noted that these criteria select for proteins significantly enriched in cilia and ciliary proteins that are found at similar levels inside and outside of cilia will be excluded from the cilia-APEX2 proteome.

**Figure 2.**
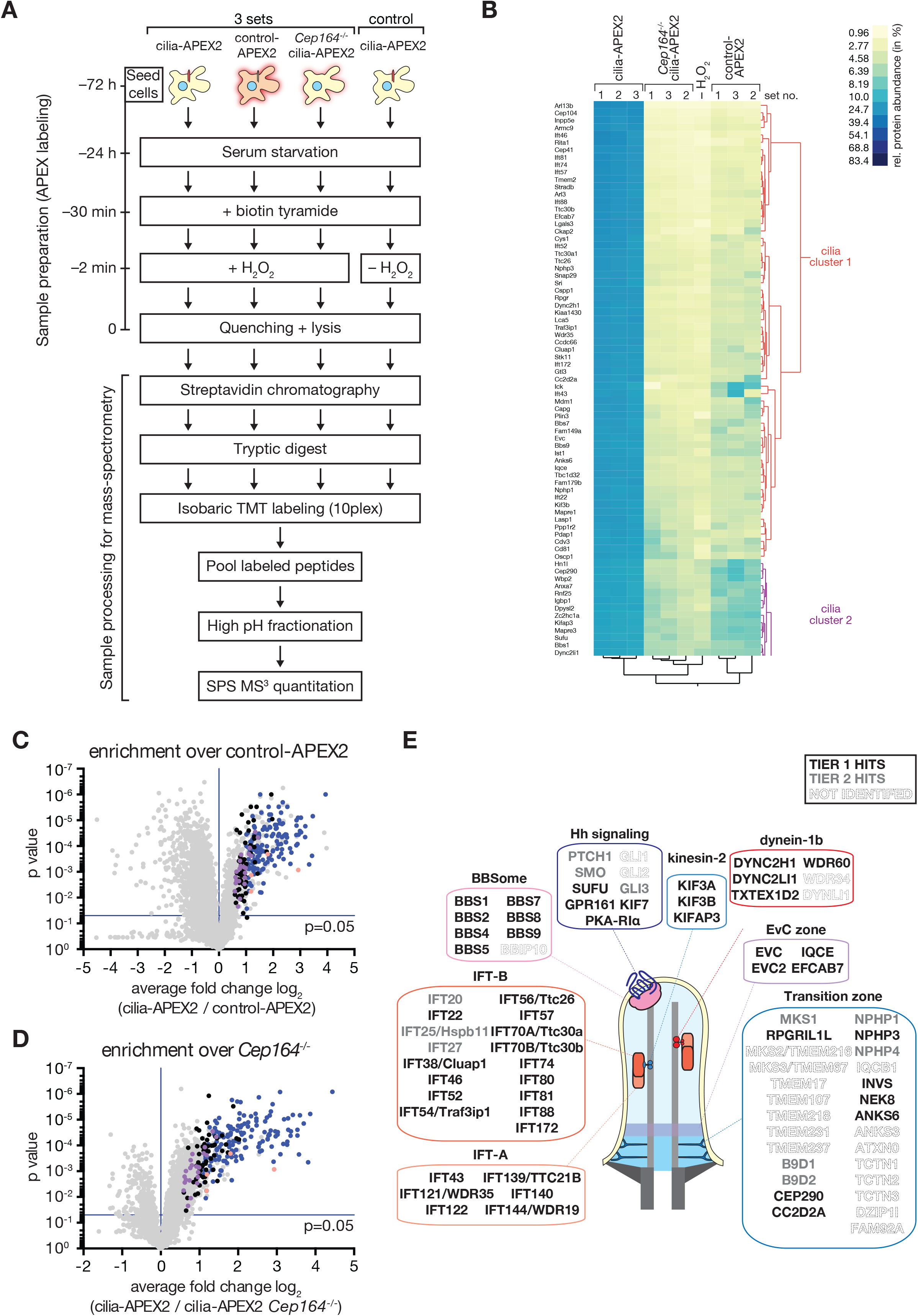
Cilia-APEX2-based proteomics reveals the proteome of primary cilia with high sensitivity. **(A)** Schematic representation of experimental workflow. Cells were seeded 72 h before the APEX labeling reaction. Cells were first grown in serum-rich medium for 48 h and then switched to serum-reduced medium to induce ciliation. APEX labeling was conducted by pre-incubating cells with biotin tyramide for 30 min and the labeling reaction was then initiated by addition of hydrogen peroxide (H_2_O_2_) for 2 min. The APEX reaction was then quenched with cyanide, ascorbate and the antioxidant Trolox and cells put on ice. In one control, H_2_O_2_ was omitted from the experimental scheme. This last control was conducted in singlet as all 10 TMT channels were then occupied. After labeling and quenching, cells were lysed and biotinylated proteins isolated by streptavidin chromatography. Bound material was extensively washed and tryptic peptides released via on-bead digest. Peptides of each individual sample were labeled with a unique tandem mass tag (TMT), all samples were pooled, and peptides were fractionated by offline high pH reversed phase chromatography to generate 12 fractions, which were analyzed using a synchronous precursor selection (SPS) MS^3^ method for mass spectrometric identification and quantitation. **(B)** Hierarchical two-way cluster analysis of a cilia-APEX2/TMT experiment conducted according to the scheme outlined in (A). Clustering of the relative abundances of each identified protein (rows) in the individual samples (columns) was performed based on Ward’s minimum variance method. The relative abundance of a given protein was calculated by dividing the TMT signal to noise ratio in one sample by the sum of TMT signal to noise ratios in all samples. Legend depicts color scheme for relative abundances (in %). The clusters containing cilia proteins are shown (see Fig. S1E for full cluster analysis). **(C** and **D)** Volcano plots showing protein enrichment in cilia-APEX2 compared to control-APEX2 samples **(C)** or in cilia-APEX2 WT vs. *Cep164^−/−^* samples **(D)**. Average enrichments of cilia-APEX2 samples versus the respective controls were plotted against the calculated *p* values (statistical significance of enrichment calculated from unpaired student’s *t*-tests). Proteins fulfilling all 4 significance and enrichment criteria are represented by blue dots, proteins that met 3 criteria are represented by black dots and proteins that failed 2 or more significance criteria are shown in grey (see text for details). Known cilia proteins outside of significance criteria or quantified from one peptide are highlighted in purple and salmon, respectively. See Table S1. **(E)** Schematic of a primary cilium with indicated hallmark protein complexes, structures or pathways. Boxes contain proteins of the respective entities identified as Tier1 (black) or Tier2 (grey) hits of the cilia-APEX2 proteome. Proteins not identified are indicated by ‘ghost’ lettering. Proteins names are shown and Gene symbols added for cases, in which gene symbols in our datasets differ from conventional protein names. Note that Ttc30a1 and Ttc30a1 have been grouped into IFT70A/Ttc30a.

To extend the coverage of the cilia-APEX2 method, we leveraged two additional independent experiments where the cilia-APEX2 *Cep164^−/−^* control was replaced by a no labeling control. We grouped the resulting candidate cilia proteins into two tiers, depending upon whether they were identified in all three datasets (Tier 1) or missing from one of the three datasets (Tier 2; Table S2). Application of these criteria resulted in the inclusion of 179 proteins in tier 1 and 91 proteins in Tier 2. While cilia-APEX had identified 75% of the IFT motor subunits, 60% of the IFT subunits and none of the BBSome subunits (Mick et al., 2015), the cilia-APEX2 proteome comprises nearly all subunits of the IFT motors kinesin-2 and dynein 2, of the IFT complexes and of the BBSome. (Fig. 2E). Most of the subunits that were not identified –e.g. BBS18 and LC8 – were less than 10 kDa and likely to be missed by MS because of the small number of tryptic peptides produced (Fig. 2E). Unexpectedly, half of the currently known components of the transition zone (TZ) were identified as hits in in the cilia-APEX2 proteome. The TZ acts as a diffusion barrier that functionally separates the cilium from the rest of the cell and was initially considered to be a static structure of the ciliary base (Garcia-Gonzalo and Reiter, 2017; Gonçalves and Pelletier, 2017). The notion of a static TZ was further reinforced by the near absence of TZ components from prior ciliary APEX studies (Mick et al., 2015; Kohli et al., 2017). However, recent studies have revealed that, while some TZ components are static, other TZ components dynamically cycle between the TZ and the cytoplasm or the ciliary shaft (Nachury and Mick, 2019) It is particularly striking to note that TZ components with previously documented dynamicity (e.g. Nphp4 and Cep290) were identified by cilia-APEX2 while TZ components that remain statically associated with the TZ over the timescale of tens of minutes (e.g. MKS2 and TMEM107) were missing from the cilia-APEX2 proteome. It is also conceivable that some TZ components identified by cilia-APEX2 are positioned at the distal side of the TZ, and hence in close proximity to the cilia-APEX2 enzyme to be labeled by biotinyl radicals. This latter hypothesis is consistent with an extension of the TZ proteins NPHP3, INVS, NEK8 into the inversin compartment that spreads from the TZ to the first few μm of the ciliary shaft.

Most importantly, cilia-APEX2 combined with TMT labeling enabled previously unachievable identification of central cilia-enriched signaling components, including most Hh signaling components known to localize to cilia (PTCH1, SMO, GPR161, KIFf7, SUFU and GLI3). The Hh transcription factor GLI1 was not identified by cilia-APEX2, consistent with undetectable GLI1 expression in the absence of Hh pathway stimulation and GLI2 could only be quantified in one (out of three) experiments, possibly because of its low abundance in IMCD3 cells.

### Time-resolved cilia-APEX2 proteomics reveals global alterations of the cilia proteome in response to Hh stimulation

Encouraged by the detection of the core Hh signaling machinery by cilia-APEX2, we sought to determine the global changes of the cilia proteome in response to Hh by subjecting cells exposed to Sonic Hedgehog (Shh) for 0, 1, 4 or 24 h to the cilia-APEX2/TMT workflow (Fig. 3A). The duplicate experimental repeats showed high reproducibility as judged by hierarchical cluster analysis of relative protein abundances (Fig. S2A). Strikingly, comparison of the cilia-APEX2 proteomes at 0 and 24 h post-Shh treatment revealed only a handful proteins with significantly changed abundance (TMT ratio > 2-fold) (Fig. 3B). Out of the 272 quantified cilia proteins (Table S2), 267 did not experience significant changes in abundance (TMT ratio < 2.0) after 24 h of Shh ligand addition. For example, the normalized protein abundance of the cilia trafficking components IFT88 or BBS1 (components of the IFT-B and BBSome protein complexes, respectively), or of other known cilia proteins, such as the inositol polyphosphate 5-phosphatase (INPP5E) or Polycystin-2 (PKD2) displayed no significant change over the course of a 24 h treatment with Shh (Fig. 4A). The low variability in relative abundances of >95% of the ciliary proteome between different time points highlights the robust and reproducible quantitation enabled by cilia-APEX2/TMT.

**Figure 3.**
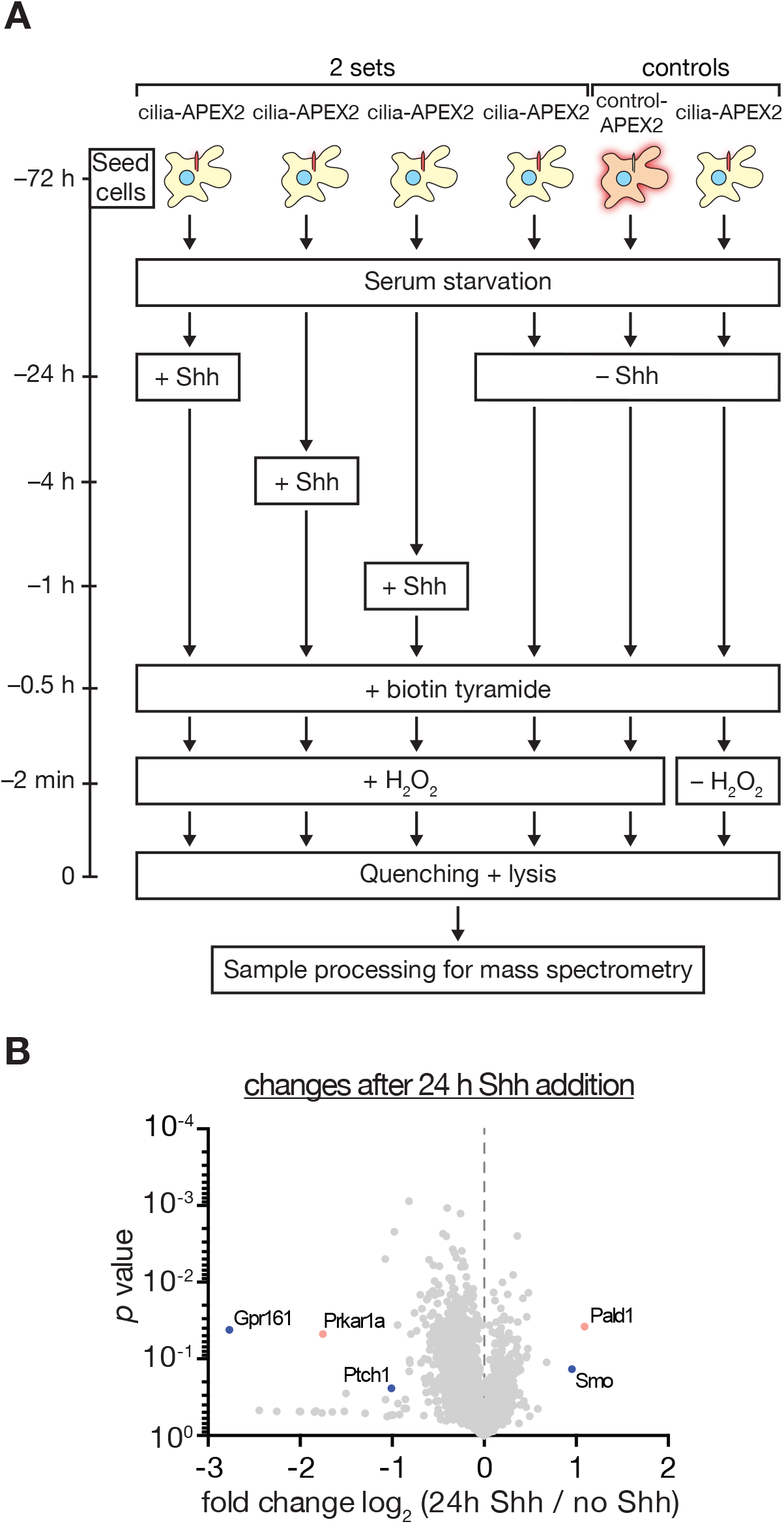
Experimental outline of time-resolved cilia-APEX2 proteomics after Sonic stimulation. **(A)** Schematic of experimental workflow for time-resolved cilia-APEX2 proteomics. Cilia-APEX2 (and control-APEX2) expressing IMCD3 cells were seeded 72 h before the APEX labeling reaction. 24 h before labeling, cells were deprived of serum, and Shh-conditioned medium was added for 24 h, 4 h or 1 h before labeling (as indicated). ‘–Shh’ indicates addition of conditioned medium without Shh. APEX labeling and sample preparation were performed as in Fig. 2A. **(B)** Volcano plot showing significance vs. enrichment in 24 h Shh-treated compared to no Shh samples. Hh signaling components known to change their ciliary localization are shown in blue, proteins with newly identified changes are shown in orange.

**Figure 4.**
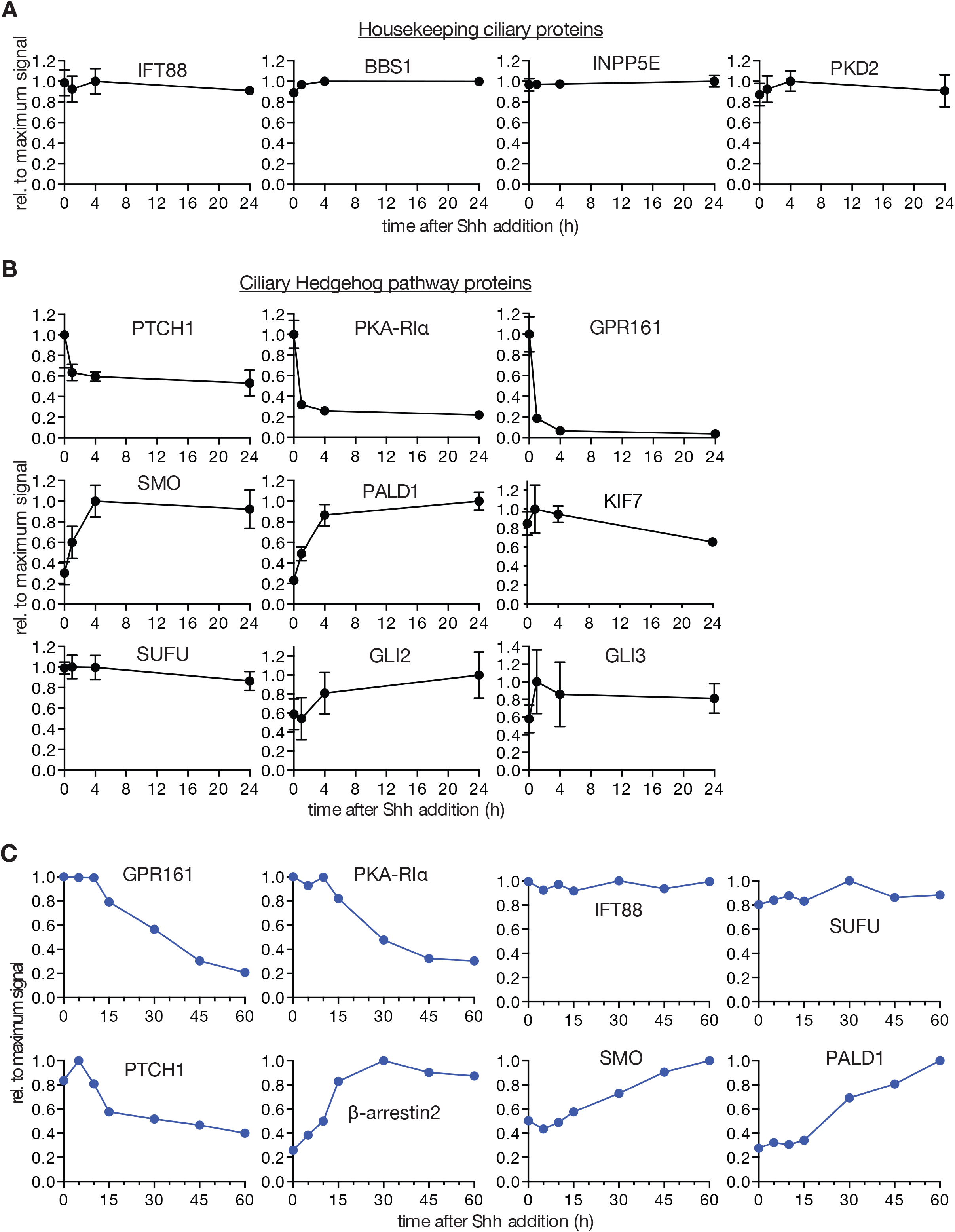
Time-resolved cilia-APEX2 proteomics reveals the extent of ciliary proteome dynamics in response to Sonic Hedgehog. **(A)** Relative protein abundances in primary cilia assessed by mass-spectrometric TMT quantitation were plotted over time. Data points represent averages of duplicate measurements, error bars depict individual values. Error bars smaller than indicated datapoint symbols have been omitted. For each individual protein, the background signal in the control-APEX2 sample was set to 0 and the maximum average signal across all time points was set to 1. 0 h, represents –Shh as in (Fig. 3A). **(B)** Hh signaling components change their abundance in cilia in response to Shh addition. Relative abundances of indicated proteins in primary cilia are plotted over time as in (**A**). **(C)** Relative abundances of indicated proteins in primary cilia were assessed by cilia-APEX2/TMT proteomics at indicated timepoints after Shh addition (see Fig. S3). Normalized intensities (relative to ARL13B) were plotted over time. Maximum signals were set to 1, TMT signals in control samples set to 0. t= 0 corresponds to the ‘–Shh’ sample.

Importantly, cilia-APEX2/TMT profiling detected changes in ciliary abundance of the three Hh signaling components known to undergo signal-dependent redistribution in or out of cilia (Fig. 4B). Levels of the Hh receptor PTCH1 and the GPCR GPR161 decreased while SMO increased within 4 h after pathway activation. The kinetics revealed by cilia-APEX2 closely matched the kinetics previously defined by immunofluorescence microscopy (Rohatgi et al., 2007; Mukhopadhyay et al., 2013). While the transcription factors GLI2 and GLI3, the GLI chaperone SUFU and the GLI regulator KIF7 become enriched at the tip of cilia in response to Hh (Wen et al., 2010; Tukachinsky et al., 2010; Endoh-Yamagami et al., 2009; Liem et al., 2009), the cilia-APEX2 signals for GLI2, GLI3 and KIF7 increased by less than 20% and the cilia-APEX2 signal for SUFU did not change during the time course of Shh treatment. We note that while GLI2 failed the stringent inclusion criteria for the cilia-APEX2 proteome, cilia-APEX2/TMT successfully quantified peptides in the 24 h time course experiment. These findings suggest that the total ciliary amounts of GLI2, GLI3, KIF7 and SUFU may not change significantly in response to Hh signal and that these factors may undergo subciliary re-distribution, *i.e.* mobilization of a broadly distributed ciliary pool to the tip. Similarly, while the BBSome becomes enriched at the tip of cilia upon Hh pathway activation (Ye et al., 2018), cilia-APEX2 signals of BBSome subunits do not change appreciably during the 24 h time course.

Since the ciliary changes of PTCH1, SMO and GPR161 were nearly complete after only 1h of pathway activation, we sought to resolve the changes in proteome remodeling during the first 60 min after Shh addition (see Fig. S3). While housekeeping ciliary proteins remained largely unchanged, the high temporal resolution and precise TMT-based quantitation of cilia-APEX2 profiling enabled a refined characterization of the redistribution of Hh signaling components (Fig. 4C). The levels of the Hh receptor PTCH1 in cilia started dropping 5 min after Shh addition and reached a minimum after 30 min. Meanwhile, ciliary levels of SMO steadily increased during the 60 min time course and until the 4 h time point (Fig. 4B). The removal of GPR161 was preceded by an increase in ciliary β-arrestin2 levels (Fig. 4C), consistent with the proposed role of β-arrestin2 in triggering signal dependent exit of GPR161 from cilia (Pal et al., 2016).

To determine if any other proteins besides SMO, PTCH1 and GPR161 undergo changes in ciliary abundance in response to Shh, we searched for proteins that co-clustered with SMO, PTCH1 or GPR161 during the 24h time course in a hierarchical two-way cluster analysis. Only one protein clustered closely with SMO (Fig. 5A), the putative phosphatase Paladin which displayed kinetics of ciliary accumulation closely mirroring those of SMO (Fig. 4B). It should be noted that despite tight clustering of PKD1 and GLI2, the PKD1/GLI2 cluster sits at a substantial distance from the SMO/PALD1 mini-cluster (Fig. 5A). Similarly, hierarchical clustering revealed only one tight cluster of proteins disappearing from cilia in response to Shh (Fig. 5B). This cluster contained GPR161, PTCH1 and the PKA regulatory subunit Iα (PKA-RIα), all of which nearly reached their minimum within the first hour of pathway induction (Fig. 4B). As observed in the 24 h time course, the ciliary exit kinetics of GPR161 and PKA-RIα were nearly identical in the 60 min time course (Fig. 4C).

**Figure 5.**
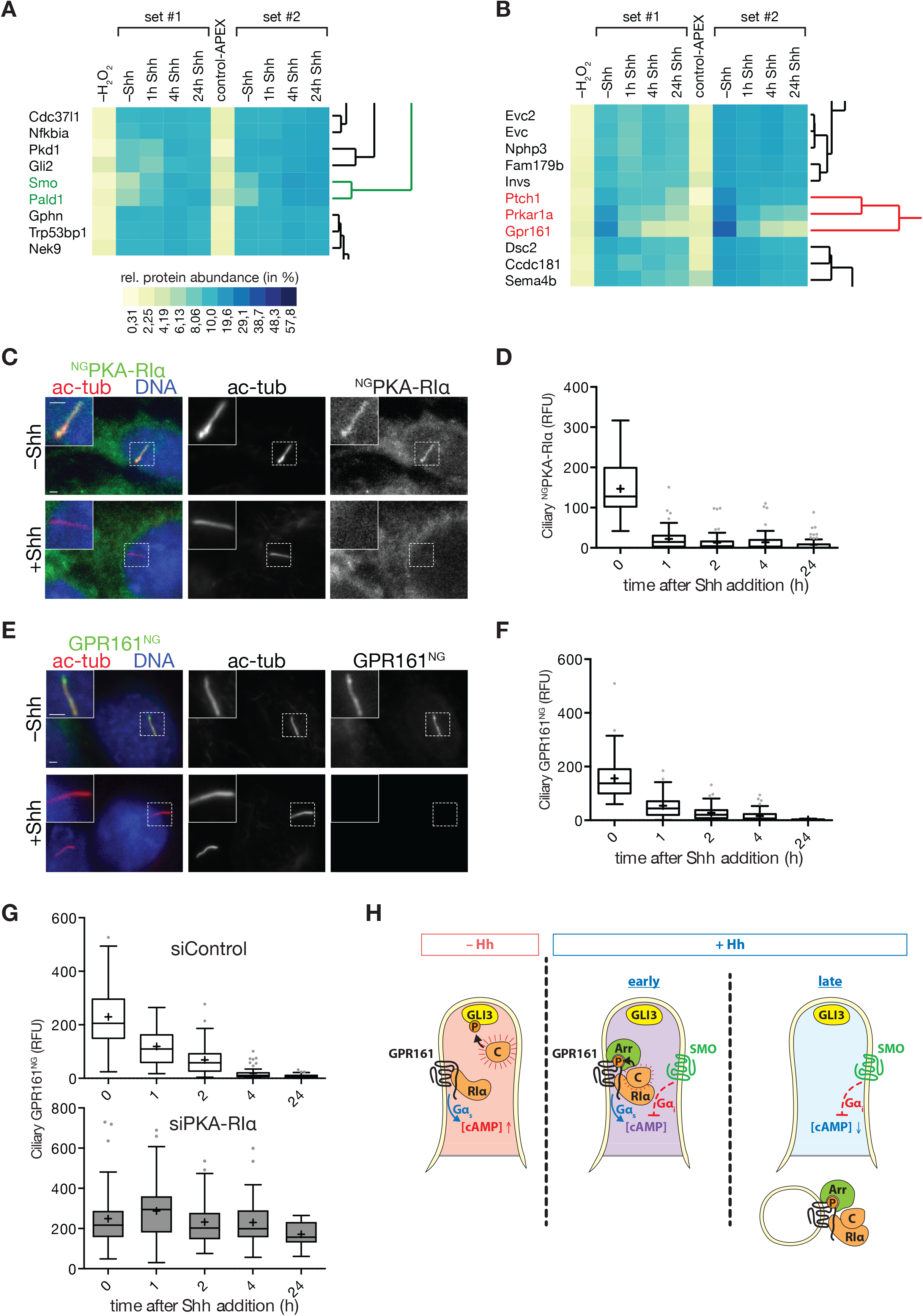
Hierarchical cluster analysis reveals ciliary exit of PKA-RIα together with GPR161 in response to Hedgehog signal. **(A)** and **(B)** Hierarchical cluster analysis of the two sets of time-resolved cilia-APEX2 proteomics experiments conducted following the scheme in Fig. 3A. (**A**) Magnified view of the SMO mini-cluster (green) and neighboring branches. (**B**) Magnified view of the GPR161 mini-cluster (red) and neighboring branches. Prkar1a is the gene name for PKA-RIα. The complete clustering analysis is shown in Fig. S2A. **(C)** IMCD3 cells stably expressing mNeonGreen-tagged PKA-RIα (^NG^PKA-RIα) were serum-starved for 24 h and treated with conditioned medium with or without Shh. Cells were fixed and stained for acetylated tubulin (ac-tub, red) and DNA (blue). ^NG^PKA-RIα was visualized by the intrinsic fluorescence of mNeonGreen (green). **(D)** Box plot showing background-corrected ^NG^PKA-RIα fluorescence in cilia at indicated timepoints after Shh addition. 60 cilia (n=<u>60</u>) were analyzed for each time point. **(E)** IMCD3 cells expressing GPR161^NG^ were treated and analyzed as in (**C**). GPR161^NG^ was visualized via the intrinsic fluorescence of mNeonGreen. **(F)** Box plot showing background corrected ciliary GPR161^NG^ signal at indicated timepoints after Shh addition. 60 cilia (n=60) were analyzed for each time point. **(G)** Box plots showing background corrected GPR161^NG^ fluorescence signals in the primary cilium of cells transfected with siRNA against *Prkar1a* or control siRNA at indicated times after Shh addition. 60 cilia were analyzed for each condition (n = 60) as in Fig. 4D. All scale bars represent 2 μm. **(H)** Model of the functional interaction between GPR161, PKA and SMO inside cilia. In unstimulated cells (–Hh), GPR161 keeps [cAMP]_cilia_ high via activation of Gα_s_. GPR161-bound PKA-RIα releases the fully active catalytic PKA subunits (C) into the lumen of cilia. Early after pathway activation (+Hh, early), SMO begins to accumulate in cilia and lowers [cAMP]_cilia_ via Gα_i_ activation. This leads to the association of the PKA-C with PKA-RIα to form a partially active holoenzyme that locally phosphorylates the GPR161 C-terminal tail. GPR161 phosphorylation results in the exit of GPR161 from cilia together with a bound PKA holoenzyme (+Hh, late). The removal of GPR161 from cilia eliminates the source of tonic Gα_s_ activation, which leads to a further reduction of [cAMP]_cilia_. ‘Arr’ indicates β-arrestin2.

Thus, cilia-APEX2/TMT not only detects known changes in remodeling of the ciliary proteome in response to Hh, the systematic nature of the TMT platform enables discovery of novel dynamic factors, co-enriched and co-depleted with known components, via kinetic profiling.

### Time-resolved cilia-APEX2/TMT illuminates the mechanisms of regulated GPR161 removal from cilia

Consistent with the current models of Hh signal transduction that predict a drop in PKA-mediated phosphorylation of GLI3 inside cilia upon Hh pathway activation, biosensor-based measurements showed that ciliary PKA activity decreases in response to Hh (Moore et al., 2016). Yet, our understanding of how ciliary PKA activity becomes depressed upon Hh pathway activation remains incomplete. On one hand, GPR161, a tonically active Gα_s_-coupled GPCR (G_s_PCR), is thought to maintain high cAMP levels in cilia of unstimulated cells. Ciliary exit of GPR161 will thus lead to a decrease of active Gα_s_ in cilia and a corresponding decrease of ciliary cAMP levels. On the other hand, the regulatory PKA subunit PKA-RIα, a stoichiometric inhibitor of the kinase-bearing catalytic PKA subunit PKA-C (Taylor et al., 2012, 201), is found inside cilia (Mick et al., 2015; Bachmann et al., 2016). PKA-RIα is thus thought to constitutively repress PKA activity inside cilia. To validate the unexpected finding that PKA-RIα may exit cilia upon Hh pathway activation, we established a stable IMCD3 cell line that expresses PKA-RIα fused to the fluorescent protein mNeonGreen (NG; (Shaner et al., 2013)) and imaged ^NG^PKA-RIα by fluorescence microscopy. While unstimulated cells exhibited robust ciliary signals of ^NG^PKA-RIα, addition of Shh triggered a decrease of ^NG^PKA-RIα ciliary fluorescence with kinetics that mirrored those measured by cilia-APEX2/TMT profiling (Figs. 5C and 5D). To pinpoint the step in the pathway that triggers the removal of PKA-RIα from cilia, we directly activated SMO via the SMO agonist SAG (Figs. S2B and S2C). The kinetics of ^NG^PKA-RIα exit from cilia were nearly identical in cells treated with Shh or SAG and we conclude that PKA-RIα exit from cilia lies downstream of SMO activation.

In agreement with our cluster analysis (Fig. 5B), the kinetics of ^NG^PKA-RIα removal from cilia upon Hh pathway activation were closely reminiscent of the exit kinetics of GPR161 (Mukhopadhyay et al., 2013), which we confirmed by imaging of GPR161^NG^ (Nager et al., 2017) (Figs. 5E, 5F, S2D). The concomitant exit of GPR161 and PKA-RIα raises the possibility that PKA-RIα piggybacks onto GPR161 to exit cilia. In support of this hypothesis, the cytoplasmic tail of GPR161 harbors an atypical A kinase anchoring protein (AKAP) motif with exquisite and unprecedented specificity for PKA-RIα (Bachmann et al., 2016). A major function of the 60 different AKAP is to recruit that the PKA regulatory subunits at discrete cellular locations to direct the catalytic subunits to their substrates (Torres-Quesada et al., 2017). We conclude that GPR161 and PKA-RIα form a stable complex that represents a functional unit, most likely together with PKA-C. Meanwhile, the abundances of other AKAPs quantified in the cilia-APEX2 dataset, e.g. AKAP11 and AKAP9, did not change appreciably in response to Hh (Fig. S2E).

A remarkable feature of the C-terminal tail of GPR161 is that it encodes both an AKAP motif for PKA-RIα and a PKA phosphorylation consensus site (Bachmann et al., 2016). In most studied instances, AKAPs interact directly with the PKA substrates (Musheshe et al., 2018) and GPR161 may further reduce the complexity of PKA recruitment to its substrate by encoding an AKAP motif within its sequence rather than interact with a separate AKAP. Because phosphomimetic mutations of the PKA site in the C-tail of GPR161 drastically reduce ciliary levels of GPR161 (Bachmann et al., 2016), we hypothesized that PKA-RIα promotes phosphorylation of GPR161 by PKA in response to Hh pathway activation and thus triggers ciliary exit of GPR161. To test this hypothesis, we assessed the Hh-induced removal of GPR161 from cilia after siRNA-mediated depletion of PKA-RIα. While control siRNA did not interfere with GPR161^NG^ exit, GPR161^NG^ failed to exit cilia in response to Hh signal in PKA-RIα-depleted cells (Fig. 5G). It is thus conceivable that the retention of GPR161 in cilia contributes to the previously reported defects in Hh pathway activation in cells depleted of PKA-RIα (Evangelista et al., 2008). A major conundrum then lies in how Hh pathway activation may control the PKA-RIα-dependent phosphorylation of the GPR161 C-tail. In the test tube, PKA regulatory subunits inhibit the activity of the catalytic subunits until the regulatory subunits bind cAMP and release free and active PKA-C. Here, recent findings that intermediate concentrations of cAMP promote PKA activation without dissociation of catalytic from regulatory subunits (Smith et al., 2017) may shed light on the regulation of GPR161 C-tail phosphorylation. While few measurements of [cAMP]_cilia_ have been published, one study found that [cAMP]_cilia_ is about 4 μM in unstimulated cells (Moore et al., 2016), a concentration sufficient to trigger nearly complete PKA-C/PKA-R dissociation within cilia. Under these circumstances, PKA-C will freely diffuse in the cilium and phosphorylate GLI2/3 and other substrates (Fig. 5H, left). Because SMO entry into cilia is already detectable at the onset of GPR161 exit (Fig. 4C), we consider the ciliary state where GPR161 and activated SMO co-exist inside cilia. We propose that the activation of Gα_i_ by SMO inside cilia will reduce [cAMP]_cilia_ to a level where an active PKA holoenzyme assembles on the C-tail of GPR161 and phosphorylates GPR161 (Fig. 5H, center). This hypothesis of ciliary Gα_i_ activation promoting phosphorylation of the C-tail of GPR161 and subsequent exit of GPR161 is supported by our previously published and yet to be explained finding that activation of the ciliary G_i_PCR SSTR3 leads the exit of GPR161 from cilia (Ye et al., 2018). The ultimate exit of the GPR161/PKA-RIα/PKA-C complex (Fig. 5H, right) further amplifies the effect of ciliary Gα_i_ activation via SMO to fully depress [cAMP]_cilia_ to a level where GLI2/3 no longer become phosphorylated by PKA. Similar to Hh ligands dually inhibiting ciliary PTCH1 by blocking its transporter activity and promoting its exit from cilia, our model proposes that PKA activity in cilia is reduced via a two-pronged mechanism that lowers cAMP levels and removes the PKA holoenzyme from cilia. Consistent with the observation that PKA-RIα is required for pathway activation in response to Smoothened agonist (Evangelista et al., 2008), our model predicts that PKA-C will remain in cilia of Hh-stimulated cells in the absence of PKA-RIα, leading to the continued production of Gli3^R^.

This model of regulated GPR161 exit resolves another conundrum raised by [Pal 2016]. Multiple groups have found that β-arrestin2 is required for exit of GPR161 from cilia subsequent to entry of activated Smoothened into cilia. Consistent with the cilia-APEX2/TMT profile of β-arrestin2 in response to Shh (Fig. 4C), recent imaging studies found that the ciliary levels of β-arrestin2 rapidly increase upon Hh pathway activation and reach a plateau at 20 min (Shinde et al., 2020). Interestingly, β-arrestin2 is also recruited to cilia upon activation of the ciliary GPCR SSTR3 and β-arrestin2 is required for removal of activated GPCRs from cilia (Green et al., 2015; Pal et al., 2016; Ye et al., 2018). Because β-arrestins are rapidly and stably recruited to activated GPCRs after they become phosphorylated, these results suggest that β-arrestin2 recognizes phosphorylated GPR161 inside cilia and instructs the ciliary export machinery to remove GPR161 from cilia. However, GPR161 was shown to be tonically active in the absence of Hh pathway stimulation (Mukhopadhyay et al., 2013). The model proposed in Fig. 5H solves this conundrum as it parsimoniously accounts for increased recruitment of β-arrestin2 to GPR161 when the entry of SMO into cilia promotes PKA-mediated phosphorylation of GPR161 C-tail. Our model also suggests that GPR161 may represent the first instance of a GPCR that is not controlled by extracellular ligands but instead monitors intracellular changes in second messenger concentrations. The GPR161/PKA-RIα module may thus act as a simple amplifier of the state of ciliary SMO activity. Considering the very recent finding that SMO can directly regulate PKA-C (Arveseth et al., 2020), our model of regulated GPR161 exit may become further refined as new details of cAMP regulation by SMO and GPR161 emerge.

While our model accounts for Hh pathway regulation by ciliary PKA-RIα, it is important to consider the extra-ciliary functions of PKA-RIα when interpreting the drastic overproduction of GLI3^R^ in unstimulated *Prkar1a^−/−^* mouse embryonic fibroblasts (Jacob et al., 2011). PKA-RIα is the only one of four PKA regulatory subunit essential for embryonic development and partial loss of PKA-RIα leads to a wide range of disease states including cystic kidneys (Veugelers et al., 2004; Amieux et al., 2002; Ye et al., 2017). Accordingly, Taylor and colleagues describe PKA-RIα as the ‘master regulator of PKA signaling’ and conclude that PKA-RIα contributes the bulk of PKA activity restriction in cells (Lu et al., 2019). Cytoplasmic PKA-C activity therefore becomes greatly elevated in the absence of PKA-RIα, and GLI2/3 phosphorylation by PKA can likely take place outside of cilia when PKA-RIα is missing. Congruent with our interpretation, overexpression of a PKA-C variant incapable of binding to the regulatory subunits potently inhibits Hh signaling, presumably by phosphorylating GLI2/3 in the cytoplasm (Iglesias-Bartolome et al., 2015).

### PALD1 accumulates within cilia of selected cell types upon Hh stimulation

Cilia-APEX2/TMT profiling detected a steady increase in SMO and PALD1 ciliary signals over the course of 60 min, and the 24 h time course indicates that SMO and PALD1 keep co-accumulating in cilia until 4 h (Figs. 4B, 4C and 5A). To validate this result, we assessed the ciliary accumulation of endogenous PALD1 by immunofluorescence microscopy (Fig. 6A). In agreement with the hierarchical clustering analysis (Fig. 5A), the increase in ciliary PALD1 signals upon Shh stimulation followed very similar kinetics to those of SMO (Fig. 6B). Given the striking Hh signal-dependent co-localization SMO and PALD1 in cilia, we sought to determine the functional interaction of PALD1 with the Hh pathway. Experimentally, activation of SMO by Smoothened agonist (SAG) fully engages the Hh pathway and bypasses PTCH1 (see Fig. 1A). Given that endogenous PALD1 and SMO accumulated efficiently in primary cilia in response to SAG (Figs. 6C-D), we conclude that ciliary accumulation of PALD1 lies downstream of SMO. Considering the nearly identical kinetics of PALD1 and SMO ciliary accumulation, we hypothesized that PALD1 may piggyback on SMO entering cilia via a physical interaction. To test this hypothesis, we leveraged cyclopamine, a SMO antagonist that promotes ciliary accumulation of SMO while blocking Hh pathway accumulation (Wilson et al., 2009; Wang et al., 2009; Rohatgi et al., 2009). While cyclopamine promoted ciliary accumulation of SMO in IMCD3 cells, it did not increase the ciliary levels of PALD1 (Figs. 6E and 6F), indicating that ciliary accumulation of SMO is not sufficient to drive PALD1 into cilia and that SMO must be activated for PALD1 to accumulate in cilia. To test whether PALD1 entry into cilia is sensitive to the reduction in [cAMP]_cilia_ downstream of SMO, we resorted to the ciliary G_i_PCR somatostatin receptor 3 (SSTR3^NG^) (Nager et al., 2017). Addition of somatostatin to IMCD3 cells stably expressing mNeonGreen-tagged SSTR3 (SSTR3^NG^) triggers activation of the GiPCR inside cilia and hallmarks of ciliary Gα_i_ activation (Ye et al., 2018). Strikingly, SSTR3 activation is sufficient to recruit PALD1 to primary cilia (Figs. 6G-H), suggesting that, similarly to GPR161 exit from cilia, ciliary PALD1 accumulation responds to drops in ciliary cAMP levels.

**Figure 6.**
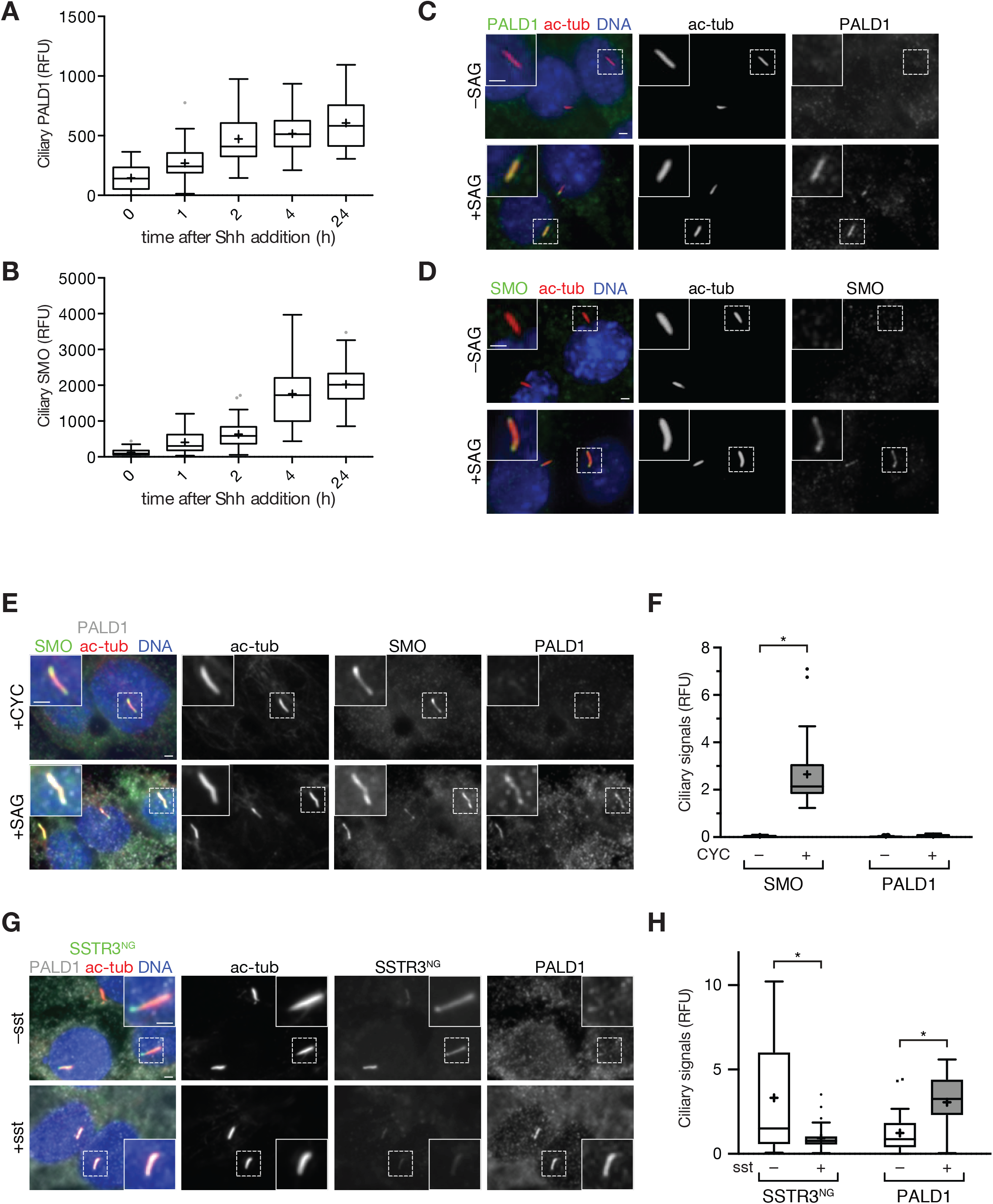
PALD1 accumulates in primary cilia in response to ciliary Gα_i_ activation. **(A** and **B)** IMCD3 cells were serum-starved for 24 h and treated with Shh for indicated times before fixation and staining for PALD1 (**A**) or SMO (**B**). Box plots display the background-corrected signals of PALD1 and SMO in primary cilia. 59 cilia were analyzed for each condition (n = 59). **(C** and **D)** IMCD3 cells were serum-starved and treated with or without SAG for 24 h and analyzed by immunofluorescence microscopy using indicated antibodies. **(E** and **F)** IMCD3 cells were serum-starved and treated with cyclopamine (+CYC) or SAG for 24 h and analyzed as in (C and D). (**E**) Micrographs of representative images. (**F**) Box plots showing background-corrected, relative ciliary fluorescence intensities of respective proteins normalized to acetylated tubulin signals. 30 cilia were analyzed for each condition (n = 30). Data was analyzed using two-way ANOVA with multiple comparisons (Tukey test) with a defined confidence of 95%. *, p < 0.05. **(G** and **H)** IMCD3 cells stably expressing Sstr3^NG^ were serum-starved for 24 h in the presence or absence of 10 μM somatostatin and analyzed as in (E and F) (n = 30). Sstr3^NG^ was detected by mNeonGreen fluorescence. Data analyzed using two-way ANOVA with multiple comparisons (Sidak test) with a defined confidence of 95%. *, p < 0.05; n.s., not significant. All scale bars represent 2 μm.

We next sought to determine whether PALD1 is generally integrated with Hh signaling, or whether PALD1 performs a more specialized function related to Hh signaling. In addition to IMCD3, PALD1 protein expression was detectable in NIH-3T3 cells, C2C12 myoblasts, MIN6 pancreatic ß cells and human embryonic kidney (HEK) cells (Fig. 7A). Unlike some Hh pathway components such as PTCH1 or GLI1 that are translational targets of the Hh pathway, PALD1 expression did not appreciably change in response to Hh pathway stimulation. Unexpectedly, PALD1 protein expression was not detectable in telomerase immortalized human retinal pigment epithelial (RPE1-hTERT) cells. Since PALD1 was detected in HEK cells, the antibody does recognize the human protein. Because RPE1-hTERT cells may not be capable of mounting a robust Hh response, we turned our attention to cell types where the Hh response has been extensively validated. NIH-3T3 cells constitute a well-accepted cell-based system for Hh signaling (Taipale et al., 2000). Surprisingly, activation of the Hh pathway in 3T3 cells led to the accumulation of SMO in cilia but failed to promote ciliary entry of PALD1 (Fig. 7B). Hedgehog signaling controls muscle differentiation (Hu et al., 2012) and the requirement for primary cilia in myoblast proliferation can be recapitulated in cultured C2C12 cells (Fu et al., 2014). While PALD1 was absent from unstimulated C2C12 cilia, PALD1 became enriched in primary cilia in response to Hh signal (Figs. 7C and 7D). These results indicate that, while not broadly generalizable, the association of PALD1 with Hedgehog signaling is not limited to IMCD3 cells.

**Figure 7.**
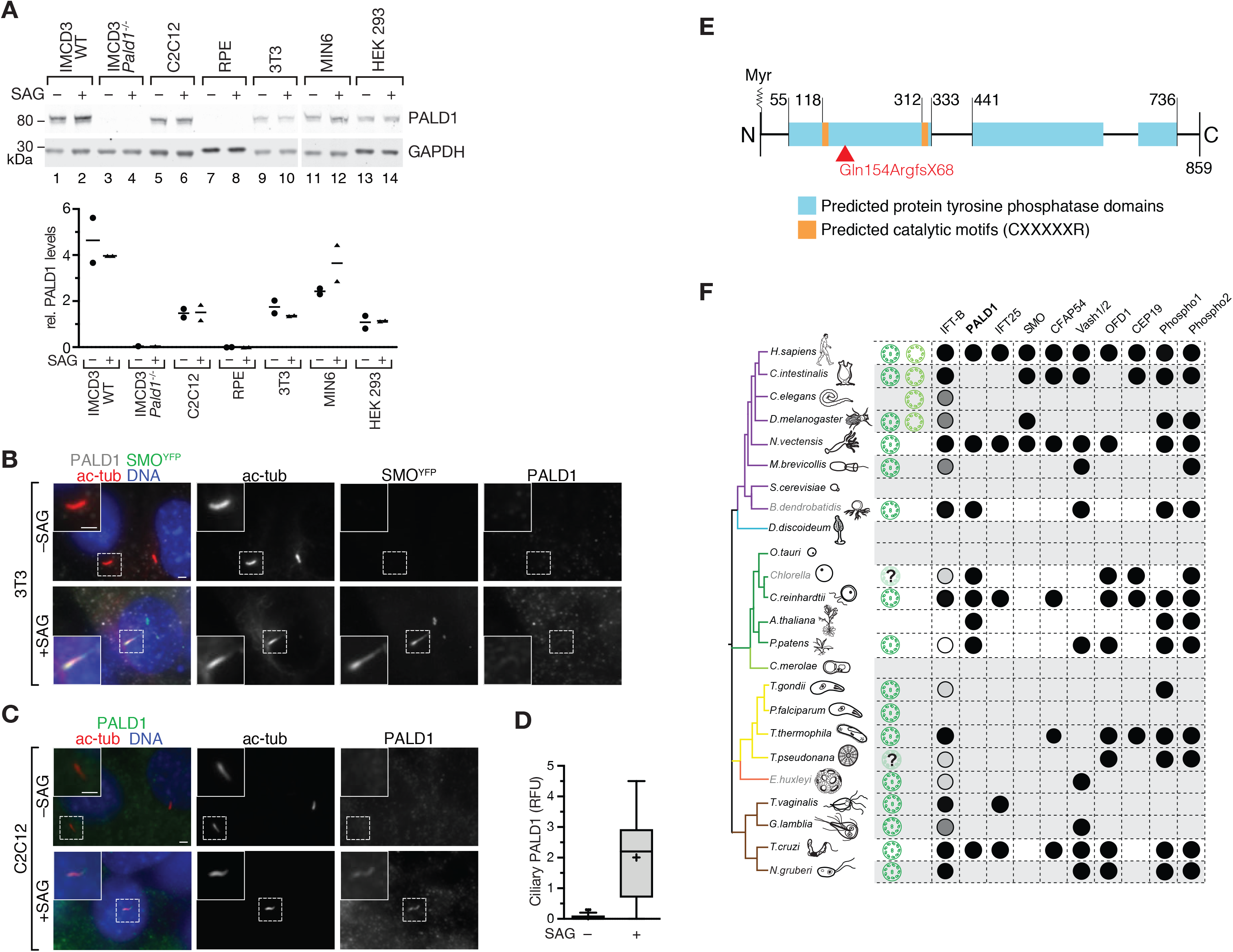
PALD1 accumulates in primary cilia of selected cell types upon Hh pathway activation. **(A)** Cell lysates of indicated cell lines were separated by SDS-PAGE and analyzed by quantitative Western blotting using anti-PALD1 antibody and anti-GAPDH as loading control. Dot plot indicates PALD1 protein levels relative to GAPDH in presence or absence of SAG as indicated (n = 2 except for for *PALD1^−/−^* where n = 1). Mean values are indicated by horizontal lines. **(B)** PALD1 does not detectably accumulate in primary cilia of 3T3 cells after Hh pathway activation whereas SMO does. 3T3 cells expressing ^YFP^SMO (Rohatgi et al., 2009) were serum-starved and treated with or without SAG for 24 h and analyzed by immunofluorescence microscopy using indicated antibodies. SMO was detected by YFP fluorescence. Scale bars = 2 μm. **(C** and **D)** PALD1 is enriched in C2C12 myoblast primary cilia after Hh pathway activation. C2C12 cells were treated and analyzed as in (B). Box plots show background-corrected, relative fluorescence normalized to acetylated tubulin signals. 30 cilia were analyzed for each condition (n = 30). **(E)** Schematic representation of PALD1 protein. Predicted protein domains and post-translational modifications. Numbers indicate amino acid positions in *Mus musculus* PALD1, Myr depicts myristoylation site at the N-terminus. Red arrow indicates location of missense mutation in *PALD1^−/−^* cells (see Fig. S4). **(F)** Phylogenetic analysis of PALD1 orthologs and co-conserved proteins (IFT25, CFAP54, Vash1/2, OFD1, CEP19, Phospho1/2) identified by Clustering by Inferred Models of Evolution (CLIME) (Li et al., 2014). The strongest co-conservation with PALD1 was observed for IFT25 (a mobile subunit of the IFT-B complex). Shown is a simplified taxonomic tree with crown eukaryotic groups in different colors (modified from (Carvalho-Santos et al., 2010)). Branch color code: purple, opisthokonts; blue, amebozoa; green, plants; yellow, alveolates and heterokonts; orange, haptophytes; and brown, excavates. When present in the respective organism, motile cilia are shown in green and primary cilia in blue. The presence of cilia in *T. pseudomonas* remains controversial. The presence of the corresponding proteins is indicated by black circles. Conservation of IFT-B complex subunits are depicted by circles with shades of grey that correspond to percentage of subunits, for which orthologs are found (black, 100%; dark grey <100%; light grey, <60%; white, <30%). The presence of orthologs was determined by CLIME, except for *B. dendrobatidis, Chlorella*, *E. huxleyi*, which were analyzed by BLASTp (blast.ncbi.nlm.nih.gov). Proteins with E-values ≤ E-25 were scored as hits.

Sequence analysis revealed several key features of PALD1 (Fig. 7E). PALD1 contains a glycine residue at amino acid position 2 that is almost certainly myristoylated as PALD1[Gly2] scores highly in all predictors of N-myristoylation (Bologna et al., 2004; Maurer-Stroh et al., 2002; Xie et al., 2016) and expression of the N-terminus of PALD1 in a cell-free system produced a protein myristoylated at Gly2 (Suzuki et al., 2010). Importantly, PALD1 was recovered in affinity purification of the myristoyl chaperone Unc119 (Wright et al., 2011). Myristoylation is a major determinant in the ciliary targeting of a variety of proteins (e.g. NPHP3, Cystin) and UNC119 mediates the entry of these myristoylated proteins into cilia (Stephen and Ismail, 2016). It is thus conceivable that PALD1’s regulated targeting to cilia involves the regulated unmasking of its attached myristate. PALD1 belongs to the large protein tyrosine phosphatase (PTP) superfamily (Chen et al., 2017). While PALD1 contains two PTP active site motifs (CXXXXXR), PALD1 is missing the 280 amino acid extended catalytic domain and represents a divergent PTP family member. In agreement with these observations, *in vitro* assays have thus far failed to detect protein phosphatase activity for PALD1 (Huang et al., 2009). In the absence of demonstrated phosphatase activity or identified substrate, PALD1 has been proposed to be an anti-phosphatase or a pseudophosphatase (Roffers-Agarwal et al., 2012).

PALD1 is conserved among all clades of eukaryotic life, from protists to mammals (Fig. 7F). Interestingly, a clustering analysis of pathways based on shared inferred ancestry (Li et al., 2014) grouped PALD1 with the cilia-associated proteins CFAP54 (cilia- and flagella-associated protein 54 (McKenzie et al., 2015, 54)), the tubulin detyrosinases vasohibin-1 and -2 (Nieuwenhuis et al., 2017; Aillaud et al., 2017), and the IFT-B subunit IFT25. Although PALD1 is not restricted to ciliated organisms, it displays some co-conservation with cilia (Fig. 7F) and the overlap in phylogenetic co-conservation is most striking with IFT25. Together with the IFT27 protein, IFT25 forms a stable subcomplex of IFT-B that functions as a regulator of BBSome function and thus participates in the regulated removal of membrane proteins from cilia (Bhogaraju et al., 2011; Liew et al., 2014; Eguether et al., 2014). Metazoa that have lost IFT25/27 either lack a Hh response (*C. elegans*) or transduce Hedgehog signals independently of cilia (*Drosophila*) (Fig. 7F). Meanwhile, organisms such as *Chlamydomonas* that have retained IFT25/27 but that do not transduce Hh signals require this IFT-B subcomplex for BBSome export from cilia (Dong et al., 2017). The shared phylogenetic pattern of association of IFT25/27 and PALD1 with Hh signaling and cilia suggests that PALD1 performs a function in cilia that supports efficient Hh signaling while not being absolutely required for either cilia assembly or Hh signaling. In agreement with the tissue-specific expression of PALD1 (Huang et al., 2009) and the integration of PALD1 with Hh signaling in a subset of cell lines, we propose that PALD1 fulfills a cell-type specific function in multicellular organisms.

Besides the aforementioned cilia-related proteins, the CLIME analysis revealed two phosphatases co-conserved with PALD1: Phospho1, an ancient phosphoethanolamine and phosphocholine phosphatase with roles in bone mineralization in vertebrates and Phospho2, a pyridoxal-5-phosphate phosphatase related to Phospho1. Phospho1/2 belong to the haloacid dehalogenase phosphatase superfamily and bear no resemblance to the PTP family or PALD1. The co-retention of Phospho1, Phospho2 and PALD1 in a variety of organism is remarkable as the only common thread between these proteins is the release of inorganic phosphate. In this context, probing the genetic interactions between PALD1, Phospho1 and Phospho2 may reveal an unexpected functional relationship between these proteins.

PALD1-deficient myoblasts exhibit increased levels of insulin receptor –a protein previously detected in primary cilia (Gerdes et al., 2014)– and PALD1 overexpression reduces insulin receptor levels as well as its signal-dependent phosphorylation (Huang et al., 2009). As we observe PALD1 accumulation in myoblast primary cilia in response to Shh but also after stimulation of the ciliary GPCR SSTR3, it is tempting to speculate that PALD1 may assist IFT25/27 in the ciliary exit of ciliary receptors and the desensitization of associated signaling pathways.

### PALD1 is an attenuator of Hh signaling in IMCD3 cells

To assess a potential role for PALD1 in Hh signaling, we generated PALD1-deficient IMCD3 cells bearing biallelic frameshift mutations by CRISPR/Cas9-mediated genome editing (Fig. S4). The morphology of cilia, as assessed by acetylated tubulin or IFT88 staining, was indistinguishable between WT and *Pald1^−/−^* cells (Fig. 8A). Immunoblotting confirmed the absence of PALD1 from *Pald1^−/−^* cells, while the levels of ciliary proteins PTCH1 and IFT88 were unaffected (Fig. 8B). Consistent with a functional relationship between PALD1 and the Hh pathway, we detected a significant reduction in the levels of GLI3 repressor (GLI3^R^) as well as an increase in full-length GLI3 (GLI3^FL^) in unstimulated *Pald1^−/−^* cells compared to WT cells (Fig. 8B). Further underscoring the similarities in phylogenetic distribution between PALD1 and IFT25/27, the effect of PALD1 deletion on GLI3 processing is reminiscent of the Hh defect described in IFT27-deficient mouse skin (Yang et al., 2015). The ratio between the full-length and repressor forms of GLI3 (GLI3^FL^/GLI3^R^) has been used as a proxy for Hh pathway activation, as this ratio increases severalfold upon Hh pathway activation (Wang et al., 2000; Wen et al., 2010). In wild-type IMCD3 cells, the GLI3^FL^/GLI3^R^ ratio increased 2-fold after Hh addition (Fig. 8C). Meanwhile, unstimulated *Pald1^−/−^* IMCD3 cells exhibit a similar GLI3^FL^/GLI3^R^ ratio as Shh-treated wild-type cells, suggesting that PALD1 restricts Hh pathway activation in the absence of Hh ligand. The GLI3^FL^/GLI3^R^ ratio further increased after Hh stimulation of *Pald1^−/−^* cells, indicating that PALD1-deficient cells still remained Hedgehog-responsive (Fig. 8C).

**Figure 8.**
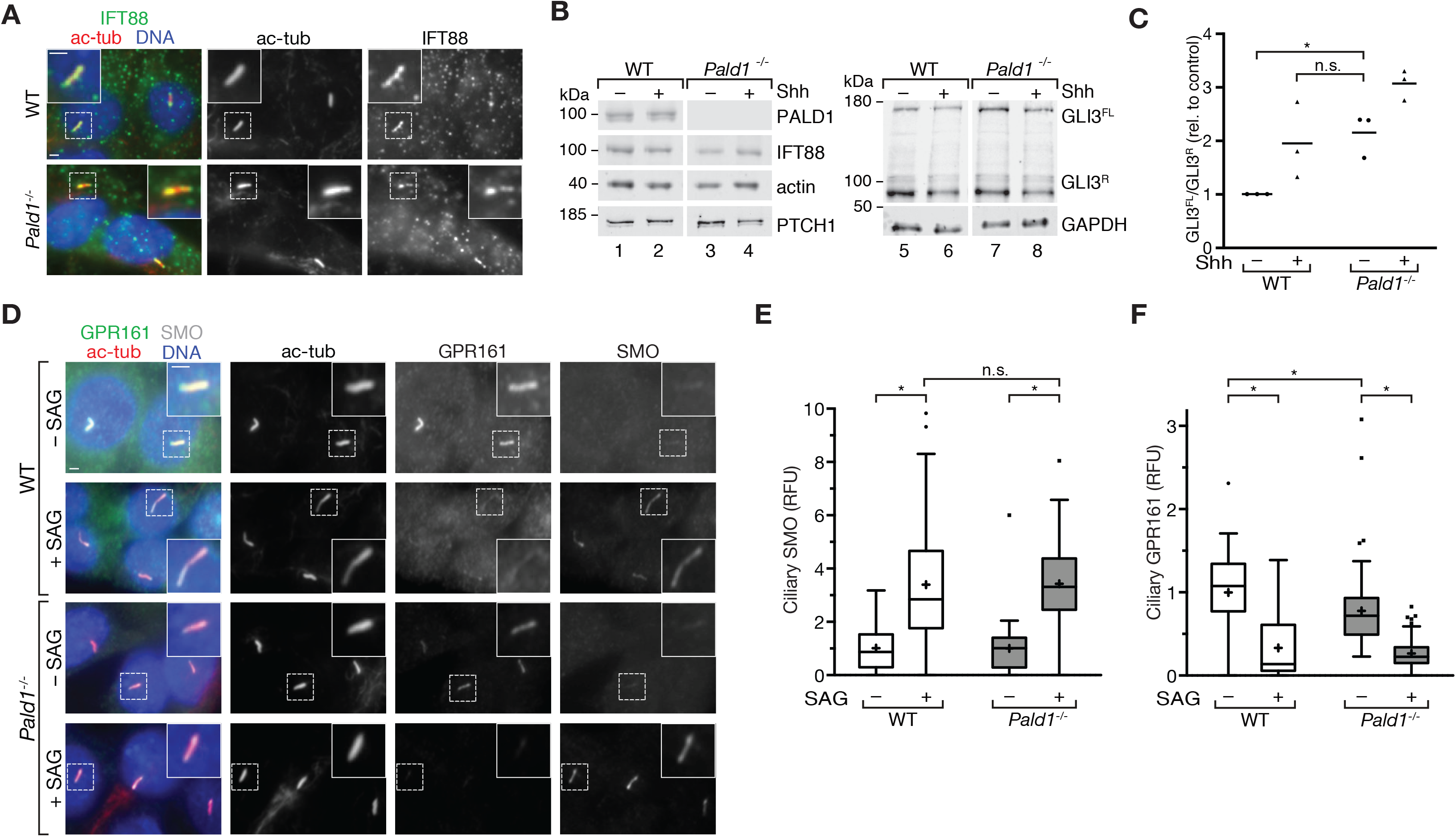
PALD1 is required for efficient GLI3 repressor formation in IMCD3 cells. **(A)** *Pald1^−/−^* and parental IMCD3 cells were serum-starved for 24 h and analyzed by immunofluorescence microscopy using indicated antibodies. **(B)** Lysates of wild-type and *Pald1^−/−^* IMCD3 cells cultured in the presence or absence of Shh were separated by SDS-PAGE and analyzed by immunoblotting using indicated antibodies. GLI3 repressor (GLI3^R^) and full-length (GLI3^FL^) forms are indicated. **(C)** GLI3^R^ and GLI3^FL^ signals from 3 independent experiments as in (B) were quantified and GLI3^FL^/GLI3^R^ ratios plotted. Horizontal lines depict means (n = 3). Each experiment was internally normalized to the GLI3^FL^/GLI3^R^ ratio in WT in the absence of signal (WT –Shh GLI3^FL^/GLI3^R^ ratio = 1). Data were analyzed using two-way ANOVA with multiple comparisons in a Tukey test with a defined confidence of 95%. *, p < 0.05; n.s., not significant. **(D)** WT and PALD1-deficient IMCD3 cells were serum-starved and treated with Smoothened agonist (SAG) for 24 h (as indicated) and analyzed as in (A). **(E)** and **(F)** Box plots showing background-corrected, relative fluorescence normalized to acetylated tubulin signals. (**E**) Two independent experiments were performed and 30 cilia were analyzed for each condition in each experiment (n = 60). (**F**) Three independent experiments were performed and 30 cilia were analyzed for each condition in each experiment (n = 90). Data were analyzed using two-way ANOVA with multiple comparisons in a Tukey test with a defined confidence of 95%. *, p < 0.05; n.s., not significant. All scale bars represent 2 μm.

To investigate the source of spontaneous pathway activation in PALD1-deficient IMCD3 cells, we assessed the cilia localization of SMO and GPR161 in response to Hh pathway stimulation. As in WT cells, SMO was not detected in resting *Pald1^−/−^* cilia and accumulated normally in response to pathway activation (Figs. 8D and 8E). Meanwhile, GPR161 levels were reduced in cilia of unstimulated PALD1-deficient IMCD3 cells compared to unstimulated WT cells (Figs. 8D and 8F), consistent with a mild spontaneous activation of the Hh pathway. In agreement with the observed increase in the GLI3^FL^/GLI3^R^ ratios, GPR161 levels in PALD1-deficient cilia decreased further after Hh pathway activation. In conclusion, PALD1-deficient IMCD3 cells do respond to Hh stimulation but spontaneously activate the pathway in IMCD3 cells. Hence, unlike for negative regulators, the Hh pathway is not strictly dependent on PALD1 function, which appears to attenuate Hh signals in certain cell types to finetune cellular responses during tissue patterning, as proposed for GPR161 (Pusapati et al., 2018a).

Numerous genetic screens identified a large number of components required for Hh signaling, including proteins required for protein trafficking and maintenance of primary cilia (Jacob et al., 2011; Breslow et al., 2018; Pusapati et al., 2018b; Kim et al., 2010; Roosing et al., 2015; Wheway et al., 2015). Here, we have identified PALD1 in our proteomic screen as a protein that accumulates in primary cilia in response to Hh signal, similar to SMO. Two aspects might explain why PALD1 had not been previously associated with Hh signaling: i) PALD1 is dispensable for Hh signal transmission in several tissues, as PALD1-deficient IMCD3 cells mount a partial response to Hh (Fig. 8) and *Pald1^−/−^* mice only show a mild phenotype (German Mouse Clinic Consortium et al., 2017). ii) PALD1 is a cell-type specific factor that accumulates in primary cilia of select cell types in response to Hh signal. The absence of signal-dependent accumulation of PALD1 in NIH-3T3 cells suggests that PALD1 may not participate in the Hh response in these cells, thus providing a possible explanation for why the functional genomics screen for Hh signaling conducted in 3T3 cells did not identify PALD1 as a regulator of Hh signaling (Breslow et al., 2018; Pusapati et al., 2018b).

Starting with the discovery that Hedgehog signaling require cilia for signal transduction in vertebrates (Huangfu et al., 2003), the past 15 years revealed an elaborate choreography of signaling factors entering and exiting cilia: one protein (SMO) that accumulates in cilia upon Hh pathway activation, two proteins (PTCH1 and GPR161) that undergo Hh-dependent exit from cilia and four proteins (GLI2, GLI3, SUFU and KIF7) that accumulate at the tip of cilia in Hh-stimulated cells. Because these findings relied on imaging the core Hh signaling components defined by *Drosophila* genetics, and since *Drosophila* transduce Hedgehog signals independently of cilia, one was left to wonder how many proteins redistributing in response to Hh were left to discover. The cilia-APEX2/TMT proteomics workflow addresses this question by enabling a systematic quantitation of individual signaling molecules in cilia in a time-resolved manner. While the discovery of novel ‘tipping’ proteins awaits the development of CiliaTip-APEX2, our analysis of the cilia proteome remodeling during Hh signaling correctly identified the Hh-dependent redistribution of SMO, PTCH1 and GPR161 and revealed two additional factors that undergo Hh-induced ciliary redistribution. PKA-RIα is likely to be part of a universal ciliary cAMP gain modulator together with GPR161 that operates in all cells, with roles possibly extending beyond Hh signaling. Interestingly, PALD1 is only connected to Hh signaling in a subset of cells. This observation highlights a critical limitation of primary cilia proteomics in mammalian cells: all available data sets are either for mouse kidney IMCD3 cells or highly specialized sensory cell types (Ishikawa et al., 2012; Mick et al., 2015; Kohli et al., 2017; Mayer et al., 2009; Liu et al., 2007; Kuhlmann et al., 2014). Given that cilia-APEX2 only relies on targeting an enzyme to cilia, it should be applicable to specific tissues and cell types, ideally from a live organism or organoids and may reveal a currently unappreciated diversity in primary cilia composition.

## ACKNOWLEDGEMENTS

We thank J. Goldstein and A. von der Malsburg for assistance in generating knockout IMCD3 cell lines, E. Krause for fluorescence-activated cell sorting, K.V. Anderson, J.K. Chen, E. Ampofo, J.M. Gerdes, C.G. Morrison, S. Mukhopadhyay, P.A. Beachy and M.P. Scott for reagents, P. Schuster for experimental assistance, D.K. Breslow, S. Schuch and S.S. Taylor for helpful discussions, and M. von Zastrow and M. Delling for comments on the manuscript. This work was supported by Deutsche Forschungsgemeinschaft (DFG) funding to D.U.M. (SFB894/TPA-22) and NIH grants to S.P.G. (GM67945) and M.V.N. (R01GM089933 and R21HD087126). This work was made possible, in part, by NEI EY002162 - Core Grant for Vision Research and by the Research to Prevent Blindness Unrestricted Grant (M.V.N.).

## AUTHOR CONTRIBUTIONS

E.A.M. performed and analyzed most experiments. D.U.M. and M.V.N. conceptualized and wrote the manuscript with support by all authors. I.G.A. conducted and analyzed the PKA-related experiments. M.K. and D.U.M. established and performed the proximity labeling, TMT labeling and mass spectrometry experiments and analyzed respective data together with S.P.G.

## AUTHOR INFORMATION

The authors declare no competing financial interests.

---

**Figure S1.**
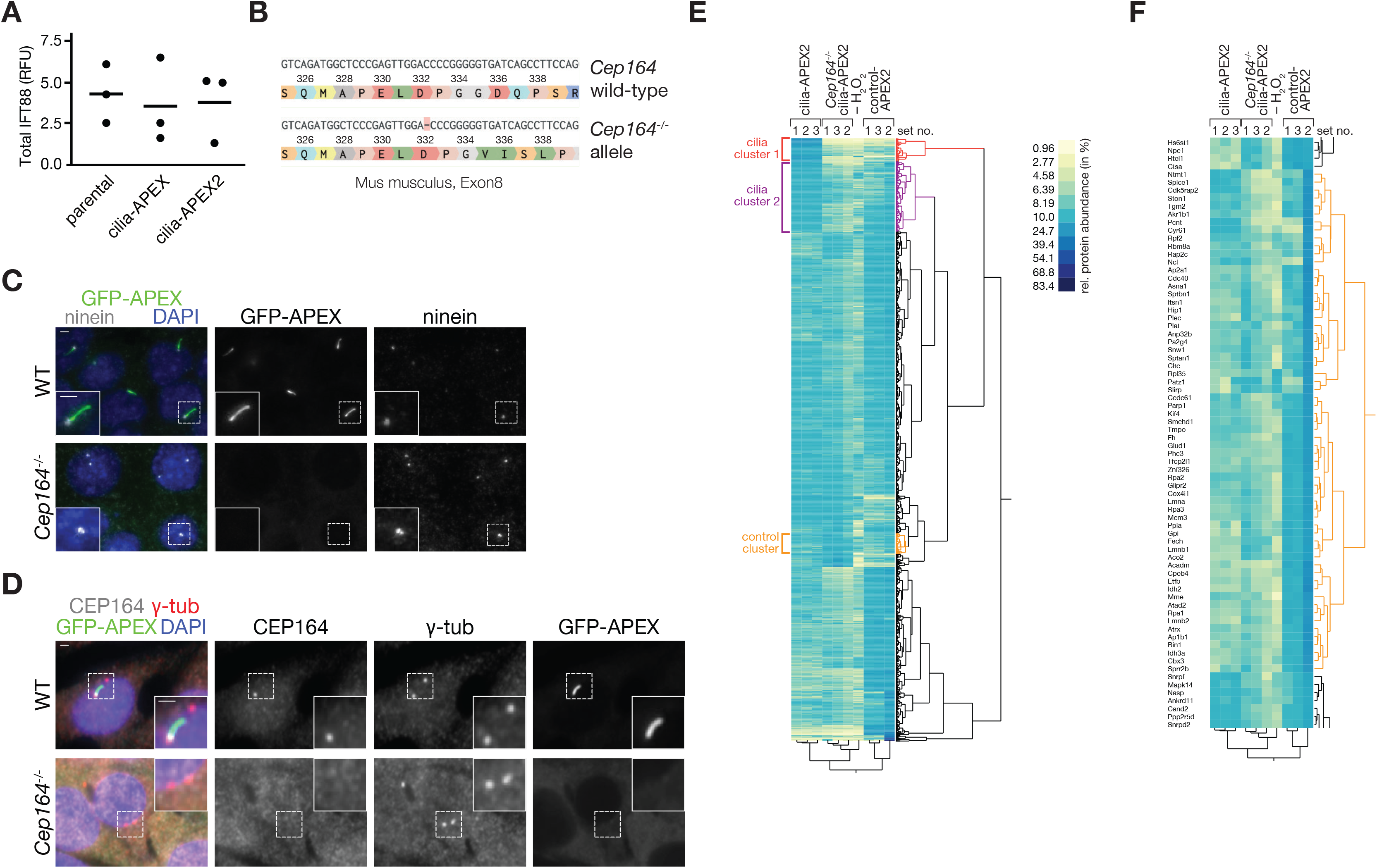
A CEP164-deficient cilia-APEX2 IMCD3 cell line serves as a specificity control for cilia-APEX2 proteomics. **(A)** Dot plot showing total protein levels of IFT88 relative to actin in indicated cell lines as determined by quantitative immunoblotting (see Fig. 1E). Mean values are indicated by horizontal lines (n = 3). **(B)** *Cep164^−/−^* cDNA was sequenced and aligned with the *Cep164* gene sequence from *Mus musculus*. A homozygous single base pair deletion in exon 8 leads to a frameshift mutation and protein truncation. DNA sequences were analyzed using Benchling. **(C)** and **(D)** Immunofluorescence micrographs of wild-type (WT) or CEP164-deficient (*Cep164^−/−^*) cell lines stably expressing cilia-APEX2. Cell lines were serum-starved for 24 h before fixation. cilia-APEX2 proteins were detected by GFP fluorescence. **(C)** Ninein marks centrioles and is visualized by antibody staining. **(D)** γ-tubulin and CEP164 are detected by specific antibodies. Representative micrographs are shown for each condition. Scale bars represent 2 μm. **(E)** High reproducibility of cilia-APEX2 proteomic setup. Hierarchical cluster analysis based on Ward’s minimum variance method (two-way clustering) of the relative abundances of each identified protein (rows) in the individual samples (columns). Relative scaled abundances were calculated by dividing the TMT signal to noise of an individual protein by the sum of TMT signal to noise ratios in all samples. Legend depicts color scheme for relative abundances (in %). Brackets indicate cilia clusters (red and purple, see Fig. 2B) and unrelated cluster (orange) without cilia proteins. **(F)** Zoom into hierarchical cluster analysis (E) at control cluster (orange) without cilia proteins.

**Figure S2.**
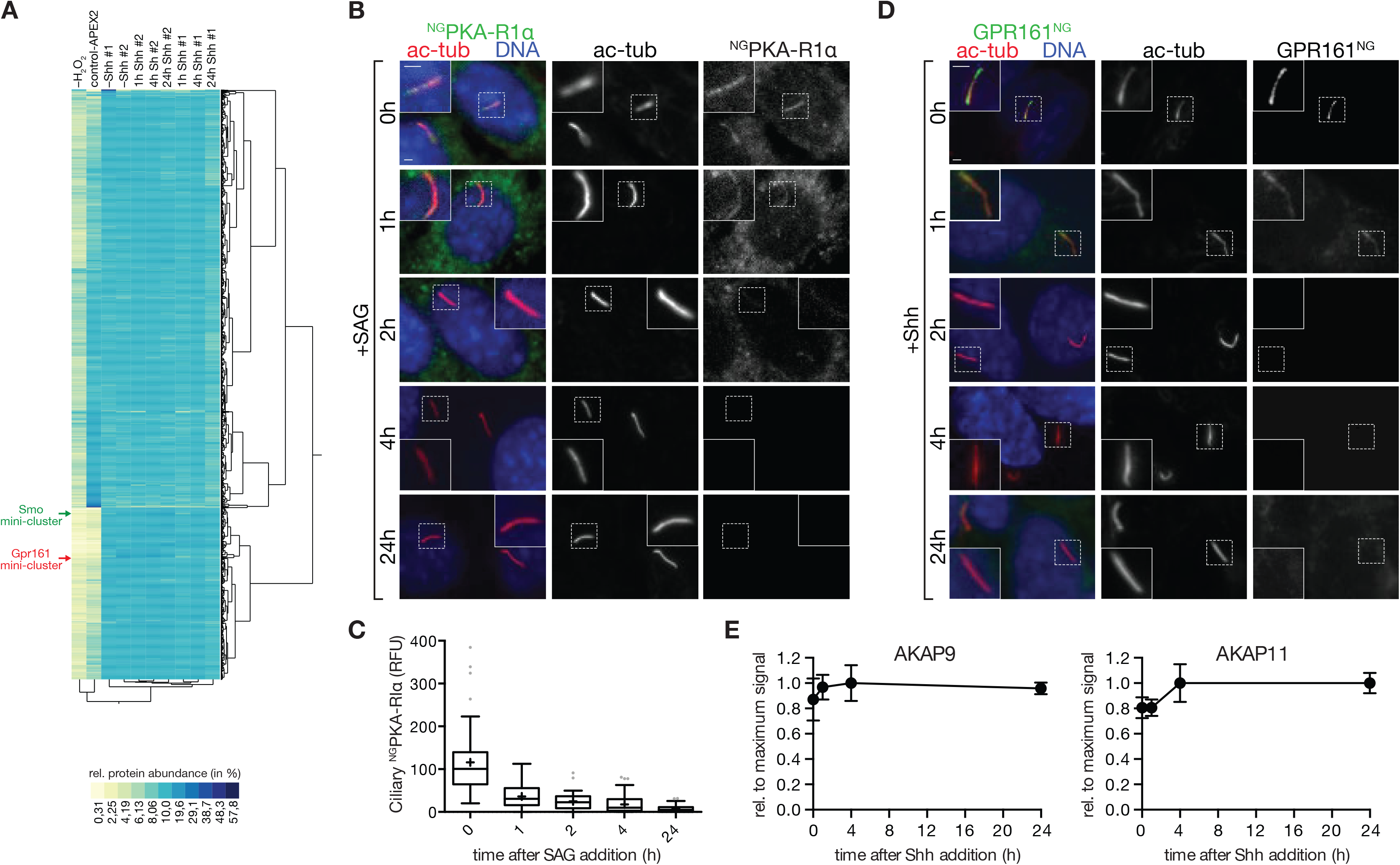
PKA-RIα and GPR161 are removed from cilia in response to Smoothened agonist, while the A-kinase anchoring proteins (AKAPs) identified by cilia-APEX2 remain unchanged. **(A)** Two-way hierarchical cluster analysis (Ward’s method) of the relative protein abundances (rows) in the individual samples (columns) shows high inter-set reproducibility. Legend depicts the color scheme of relative abundances (calculated as in Fig. S1E). SMO and GPR161 mini-clusters are indicated by green and red arrows, respectively (see Fig. 5A and B for magnified views of the mini-clusters). **(B** and **C)** ^NG^PKA-RIα-expressing IMCD3 cells were serum-starved for 24 h and treated with or without SAG for indicated times. Cells were fixed and stained for ac-tub (red) and DNA (blue). ^NG^PKA-RIα was visualized by mNeonGreen fluorescence. Representative micrographs are shown (**B**). (**C**) Box plot shows background-corrected ^NG^PKA-RIα fluorescence in cilia at indicated timepoints after SAG addition. 50 cilia (n=50) were analyzed for each time point. **(D)** IMCD3 cells expressing GPR161^NG^ were serum-starved for 24 h in the presence of Shh-conditioned medium for indicated times and analyzed as in (**B**). GPR161^NG^ was detected by mNeonGreen fluorescence. For quantitative analysis, see Fig. 5F. **(E)** Relative AKAP abundances assessed by mass-spectrometric TMT quantitation were plotted over time. Data points represent averages of duplicate measurements, error bars depict individual values. Maximum average signal was set to 1, background signals as assessed from control-APEX2 labeled samples set to 0. ‘0 h’ represents –Shh. Scale bars represent 2 μm.

**Figure S3.**
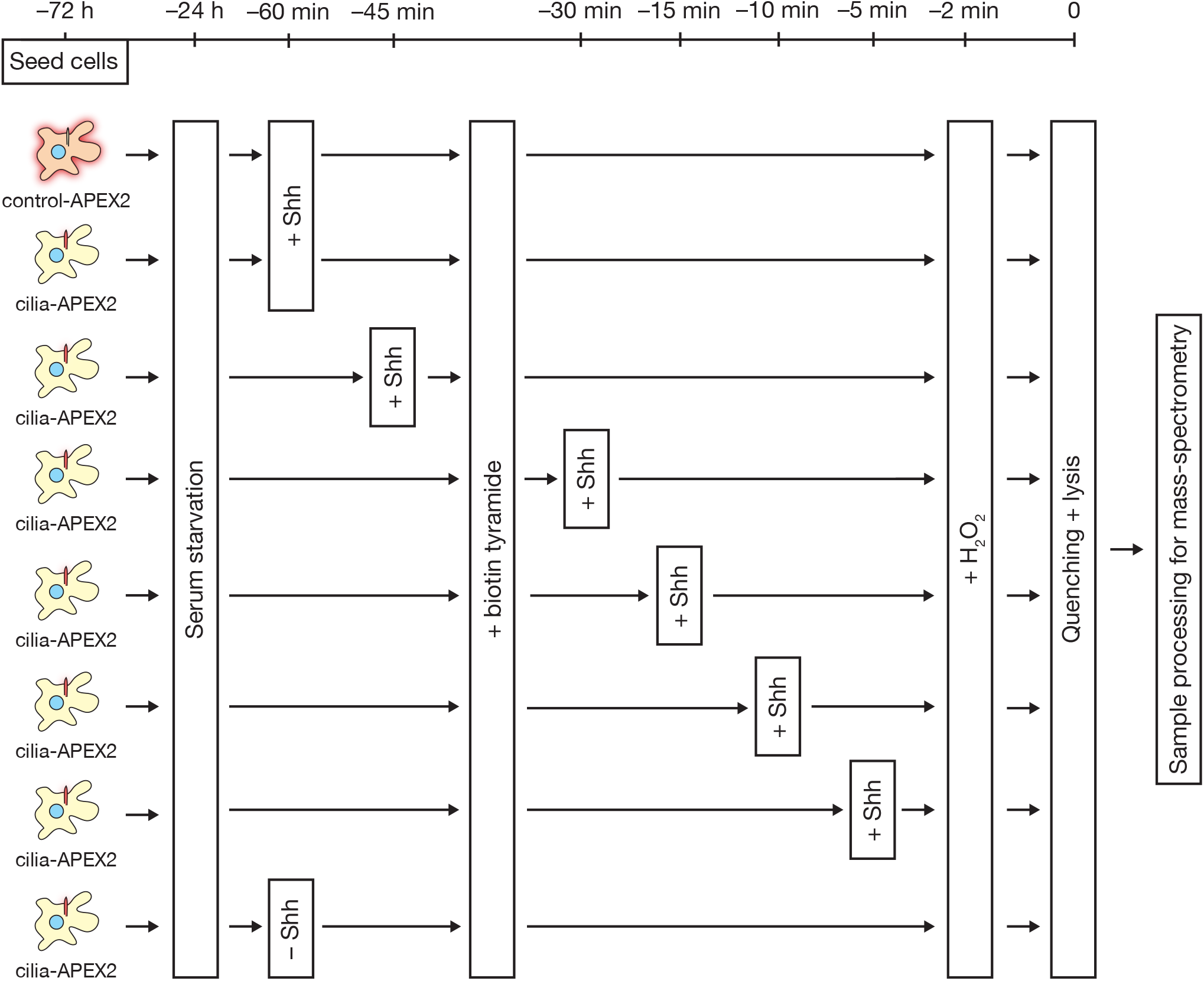
Schematic of experimental workflow for high resolution time-resolved cilia-APEX2 profiling of the ciliary Hh response (see Figure 4C). Cilia-APEX2 (and control-APEX2) IMCD3 cells were seeded 72 h before the APEX labeling reaction. 24 h before labeling, cells were starved of serum. 60 min, 45 min, 30 min, 15 min, 10 min and 5 min before APEX-labeling Shh-conditioned medium was added (as indicated). ‘–Shh’ indicates addition of conditioned medium without Shh 60 min prior to labeling. APEX labeling and sample preparation were performed as in Fig. 2A. In brief, biotin tyramide was added 30 min before the 2 min labeling reaction in the presence of hydrogen peroxides (H_2_O_2_). Samples were quenched and kept on ice for lysis, followed by streptavidin chromatography and on-bead tryptic digestion. Peptides of each sample were labeled with individual tandem-mass-tags (TMTs), pooled and fractionated offline via high pH reversed phase chromatography before mass-spectrometric analysis.

**Figure S4.**
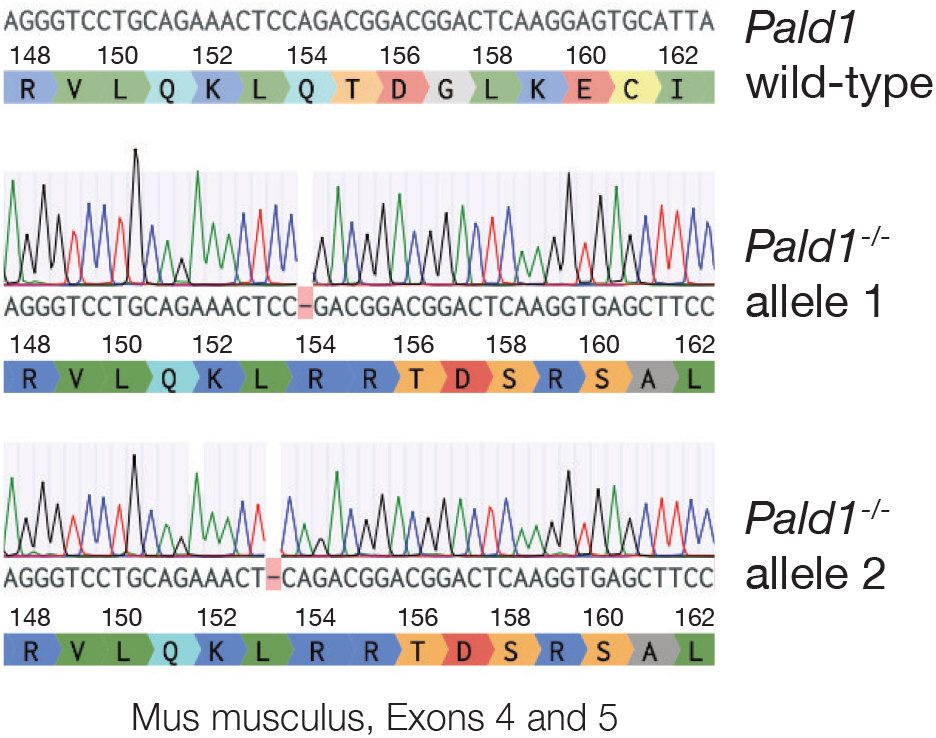
*Pald1^−/−^* IMCD3 cells exhibit early missense mutation. *Pald1^−/−^* genomic DNA was sequenced and aligned with the *PALD1* WT gene sequence from *Mus musculus*. Single base pair deletions in exon 4 lead to frameshift mutations in both alleles and protein truncation. The DNA sequences have been analyzed using Benchling.

## SUPPLEMENTARY TABLES

Table S1. **Cilia-APEX2/TMT proteomics reveals the proteome of primary cilia with high sensitivity.**

First tab ‘Legend’ shows the color scheme. Second tab ‘cilia-APEX2 TMT - Fig. 2’ enumerates the candidate cilia proteins identified by cilia-APEX2/TMT proteomics (see Figs. 2A-D). Proteins are listed by Gene symbols according to the *Mus musculus* proteome database (Uniprot 07/2014). Column B shows the numbers of quantified peptides. ‘TMT ratio’ in columns C and E is the average relative enrichment between three cilia-APEX2 samples and the indicated control samples. *p* values in columns D and F were calculated from multiple *t*-tests (statistical significance of enrichments of cilia-APEX2 samples versus the respective controls determined by unpaired student’s *t*-tests).Third tab ‘known cilia proteins below cut’ lists known cilia proteins quantified by cilia-APEX2/TMT proteomics that did not meet the inclusion criteria.

Table S2. **Cilia-APEX2 proteome list.**

First tab ‘Legend’ shows the color scheme. Second tab ‘cilia-APEX2 proteome’ lists the high confidence cilia proteins that have been scored as hits in three (Tier 1, detailed in third tab) or at least two out of three experiments (Tier 2, detailed in fourth tab). The numbers of quantified peptides and the average enrichment scores (TMT ratios) in the individual experiments are shown.

## MATERIALS AND METHODS

### Cell line generation, cultivation and manipulation

C2C12 myoblasts, NIH 3T3 and HEK cells were cultured in DMEM medium, RPE1-hTERT (described as RPE in the text) and all IMCD3 cell lines were grown in DMEM/F12, all supplemented with 10% FBS. The pancreatic beta cell line MIN6 was cultured in DMEM medium, supplemented with 15 % FBS. Ciliation was induced by growth factor deprivation by reducing the growth media to 0.2% FBS for 24 h. Transfections were carried out using XtremeGene9 (Roche) or FuGene6 (Promega) according to manufacturers’ guidelines. *Cep164* and *Pald1* genes were disrupted in IMCD3 FlpIn cells using CRISPR/Cas9-mediated genome editing with gRNAs targeting exons 8 and 4, respectively (Ran et al., 2013). Clones of each cell line were obtained by limited dilution (*Cep164*) or single cell sorting (*Pald1)*. Clones with disrupted genes were screened by immunofluorescence microscopy and Western Blotting using protein-specific antibodies. Selected positive clones were further characterized by sequencing, confirming missense mutation leading to early termination of translation.

IMCD3 cell lines stably expressing cilia-APEX2, control-APEX2, ^NG^PKA-RIα were generated using the FlpIn system as described (Breslow and Nachury, 2015). Plasmids encoding cilia-APEX2 and control-APEX2 were created by site-directed mutagenesis of cilia-APEX and control-APEX plasmids, and confirmed by sequencing. A plasmid encoding ^NG^PKA-RIα was generated using the Gateway cloning system (Life technologies) by LR clonase reaction of pEF5/FRT/NG-DEST with pENTR-PRKAR1A (obtained from Addgene, #23741). IMCD3 cells stably expressing GPR161^NG^ have previously been described (Ye et al., 2018).

To induce Hh signaling, growth media were supplemented with either 200 nM Smoothened agonist (SAG) or Shh-N conditioned medium (10-16% (v/v) depending on batch) produced with EcR-ShhN cells (gift from Phil Beachy). To block Hh signaling, cyclopamine (CYC) was added to the growth medium to a final concentration of 10 μM.

### APEX labeling experiments

Cells were incubated in the presence of 0.5 mM biotin tyramide for 30 min before the addition of hydrogen peroxide (H_2_O_2_) to a final concentration of 1 mM. For non-labeling samples water was added instead of H_2_O_2_. After 2 min of incubation at room temperature, the medium was aspirated quickly and cells were washed three times with quenching buffer (1x PBS supplemented with 10 mM sodium ascorbate, 10 mM sodium azide and 5 mM Trolox). For fluorescence microscopy, cells were immediately fixed. For proteomic and Western Blot analyses cells were lysed by scraping them off the growth surface in ice-cold lysis buffer (0.5 % (v/v) Triton X-100, 0.1 % (w/v) SDS, 10% (w/v) glycerol, 300 mM NaCl, 100 mM Tris/HCl pH 7.5, protease inhibitors) supplemented with 10 mM sodium ascorbate, 10 mM sodium azide and 5 mM Trolox. After collecting the lysate in a reaction tube, the lysate was vortexed, incubated on ice for 15 min and cleared by centrifugation (16,000 x g for 20 min at 4°C).

### Streptavidin chromatography

After determining protein concentrations of lysates from APEX-labeling experiments they were adjusted to equal concentrations and volumes as starting material for chromatography, from which samples were taken as loading control for SDS-PAGE and Western Blot analysis. Samples were added onto washed and equilibrated Streptavidin-Sepharose beads (Thermo Scientific) and biotinylated proteins were allowed to bind for 1 h at room temperature. Unbound material was removed and samples taken for Western Blot analysis. Beads with bound proteins were washed extensively with lysis buffer, then with urea wash buffer (4 M urea 10 mM Tris/HCl, pH 7.5) and finally with urea wash buffer supplemented with 50 μM biotin. For mass-spectrometric analyses, bound proteins were alkylated and digested with endopeptidase Lys-C (Wako) for 3 hours and trypsin (Promega) on beads overnight at 37°C.

### Mass spectrometry

Tryptic digests were directly labelled in 200 mM HEPES pH 8.5 with tandem mass tag (TMT) 10-plex reagents (Thermo Fisher Scientific #90406). After efficient labeling was checked by MS, peptides were subjected to alkaline reversed phase fractionation as described (Paek et al., 2017). Pooled fractions were analyzed on a Fusion Lumos Orbitrap mass spectrometer coupled to a Proxeon EASY-nLC 1000 liquid chromatography (LC) system (Thermo Fisher Scientific) using a synchronous precursor selection (SPS) MS^3^ method (McAllister and Gygi, 2013). Capillary columns had an inner diameter of 100 ☐m and were packed with 2.6 ☐m Accucore beads (Thermo Fisher Scientific). Peptides were analyzed on acidic acetonitrile gradients for 5 h with MS^1^ (Orbitrap, resolution 120k) scans, MS^2^ scans after collision-induced dissociation (CID, CE-35) in the ion trap and MS^3^ precursor fragmentation by high-energy collision-induced dissociation (HCD). Reporter ions were analyzed by MS^3^ in the orbitrap at a resolution of 50k. Further details on LC and MS parameters can be found in (Paek et al., 2017).

### Mass spectrometry data analysis

Mass spectra were processed and peptide-spectrum matches (PSM) were obtained by a SEQUEST (V.28, rev. 12) based software. Searches used a size-sorted forward and reverse database of the *M. musculus* proteome (Uniprot 07/2014) with a mass tolerance of 20 ppm for precursors and a fragment ion tolerance of 0.9 Da. Oxidized methionine residues were dynamically searched (+15.9949 Da). A false discovery rate of 1% was set for PSM following linear discriminant analysis and FDR for final collapsed proteins was 1% as well.

Relative protein quantification used summed MS^3^ TMT signal / noise (s/n) per protein filtered for summed s/n >180 over all channels per peptide and an isolation specificity >70% for each peptide. Details of the TMT intensity quantification method can be found in (Paulo et al., 2016).

### Immunofluorescence microscopy

For microscopic analyses all cells were grown on round 12 mm #1.5 coverslips and fixed in 4% paraformaldehyde for 10 to 15 min at room temperature. After fixation cells were permeabilized in −20°C methanol for 5 min and rehydrated in 1x PBS at room temperature. After extensive washing in 1x PBS, fixed cells were blocked in blocking buffer (3% bovine serum albumin, 5% serum in 1x PBS) for 30 min. After blocking, cells were incubated with primary antibody dilutions in blocking buffer for 1 h at room temperature or 4°C overnight, washed three times with 1x PBS over 15 minutes, and incubated with AlexaFluor488-, Cy3- or Cy5-coupled secondary antibodies (Jackson Immunoresearch), or AlexaFluor647-conjugated streptavidin (Invitrogen) in blocking buffer for 30 min. Finally, cells were washed five times in PBS and mounted on glass slides using Roti®-Mount FluorCare DAPI (Carl Roth; Figs. 1, 6C, 6D, 7 and 8) or DNA stained with Hoechst 33258 and mounted on glass slides using Fluoromount G (Electron Microscopy Sciences; Figs. 5, 6E, 6G, S1). APEX enzymes and ^YFP^SMO were detected by GFP and YFP fluorescence, ^NG^PKA-RIα and GPR161^NG^ by mNeonGreen fluorescence.

Prepared specimens were imaged on an AxioImager.M1 microscope (Carl Zeiss, Figs. 5C, 5E, 5G) or a Leica DMi8 with PlanApochromat oil objectives (63x, 1.4NA) using appropriate filters. Images were captured using a CoolSNAP HQ (Photometrics) or Leica DFC3000 G camera system, respectively.

### SDS-PAGE and Western Blotting

Standard techniques were used for SDS-PAGE and Western blotting. Cell lysates were generated after washing cells with 1x PBS and scraping cells of the growth surface in solubilization buffer (25 mM Tris/HCl, 300 mM NaCl, 1 mM EDTA, 10 % Glycerol, 1 % Triton X-100 (v/v), 0.1 % SDS (w/v), 1 mM PMSF and proteinase inhibitors (Roche)). Lysates were cleared by centrifugation (20.000 g at 4°C for 45 min), 25 μg of protein was separated on 10% Bis-Tris polyacrylamide gels and transferred onto nitrocellulose membranes. After blocking in 5% milk or Intercept® (TBS) Protein-Free Blocking Buffer (LI-COR) and specific antibody decoration, membranes were washed and primary antibodies visualized using IRDye800-conjugated and IRDye680-conjugated secondary antibodies on a LI-COR Odyssey laser scanner. Quantitation of bands was performed using the Image Studio Lite software (version 5.2.5).

### Statistical analyses

Statistical analyses were performed with Graphpad Prism v8.3.1. For Western Blot analyses mean values from independent experiments (exact n stated in figure legend) were calculated and are shown with either SD or SEM as described in the figure legends. For GLI3 analysis, each experiment was internally normalized to the GLI3^FL^/GLI3^R^ ratio in WT cells in the absence of Shh-N. For Immunofluorescence experiments at least 30 cilia were analyzed (exact n stated in figure legends). To compare biological replicates, relative fluorescence values were normalized to the average relative fluorescence signal in WT cells in the absence of inducing reagents. In all statistically analyzed experiments, significance was assessed by two-way ANOVA assuming normal distribution and multiplicity adjusted p-values were obtained by Holm-Sidak post-hoc or Tukey testing (p<0.05 was considered statistically significant). For volcano plot graphs, Student’s t-test were used and statistical analyses were performed using Prism 8 (Graphpad Software). Hierarchical cluster analyses were performed according to Ward’s minimum variance method using JMP software (from Statistical Analysis System, v15.1.0).

### Antibodies and reagents

Antibodies against the following proteins were used at indicated dilutions: anti-acTub (Sigma-Aldrich T7451, mouse 1:2,000), anti-Arl13b (Proteintech 17711-1-AP, rabbit 1:2,000), anti-IFT88 (Proteintech 13967-1-AP, rabbit 1:200 in IF, 1:1,000 in WB), anti-GFP (raised against 6His-tagged eGFP, rabbit 1:1,000), anti-GPR161 (gift from S. Mukhopadhyay, rabbit 1:500), anti-GLI3 (R&D Systems nachuryAF3690, goat 1:1,000) anti-actin (self-made, rabbit 1:5,000), anti-GAPDH (Proteintech 60004-1-Ig, mouse 1:2,000), anti-PALD1 (Sigma-Aldrich HPA017343, rabbit, IF: 1:250, WB: 1:1,000), anti-SMO (Santa Cruz sc-166685, mouse IgG2a in IF 1:200; Abcam ab236465 for Figs. 5B, D and E), anti-PTCH1 (Abcam ab53715, rabbit, WB 1:1,000), anti-ninein (gift from M. Bornens, rabbit 1:10,000), anti-CEP164 (gift from C. Morison, rabbit, IF 1:2,000), γ-tubulin (Proteintech 66320-1-AP, rabbit 1:1,000). Streptavidin-pHRP (Thermo Scientific #21140, 1:1,000), Streptavidin-AF647 (Invitrogen S21374, 1:1,000), Smoothened agonist (SAG; Abcam ab142160), Shh (N-terminus of Shh, produced in cell line EcR-ShhN cells, gift from Phil Beachy), cyclopamine (CYC; Merck Millipore #239806), biotin tyramide (Iris Biotech GmbH), somatostatin (sst; Alfa Aesar J66168).

### Cell lines used in this study

| Cell line | Parental cell line | reference |
| --- | --- | --- |
| *Cep164*<sup>-/-</sup> | IMCD3 FlpIn | This work |
| *Pald1*<sup>-/-</sup> | IMCD3 FlpIn | This work |
| Cilia-APEX2 | IMCD3 FlpIn | This work |
| Control-APEX2 | IMCD3 FlpIn | This work |
| *Cep164*<sup>-/-</sup> cilia-APEX2 | IMCD3 FlpIn | This work |
| <sup>NG</sup>PKA-RI $\alpha$ | IMCD3 FlpIn | This work |
| GPR161<sup>NG</sup> | IMCD3 FlpIn | (Nager et al., 2017) |
| SSTR3<sup>NG</sup> | IMCD3 FlpIn | (Nager et al., 2017) |
| <sup>YFP</sup>SMO | 3T3 | (Milenkovic et al., 2009) |
| C2C12 | - | (Blau et al., 1985) |
| RPE1-hTERT | - | (Bodnar et al., 1998) |
| MIN6 | - | (Poitout et al., 1995) |

